# *Dnmt1*-deficiency in PV interneurons alters cortical circuit function and leads to depression-like behavior

**DOI:** 10.1101/2024.09.28.615593

**Authors:** Jenice Linde, Gerion Nabbefeld, Severin Graff, Can Bora Yildiz, Daniel Pensold, Julia Reichard, Marie Hermanns, Björn Kampa, Anja Urbach, Simon Musall, Geraldine Zimmer-Bensch

## Abstract

Neuropsychiatric disorders, including major depressive disorder (MDD), are highly prevalent in modern society, arising from a complex interplay of genetic and environmental factors. Alterations in the function of cortical inhibitory GABAergic interneurons, along with dysregulations of epigenetic signatures and key regulators such as DNA methyltransferase 1 (DNMT1), have been implicated in these conditions. Through its role in catalyzing DNA methylation, DNMT1 modulates the synaptic activity of parvalbumin-expressing (PV) interneurons, which are essential for cortical inhibition. However, the functional consequences of DNMT1 activity in cortical interneurons at the network level and its impact on behavior remain unknown and must be explored to fully understand the disease implications of dysregulated DNMT1 expression and function. To address this, we utilized a conditional knockout mouse model with *Dnmt1* deletion in PV interneurons. Our findings reveal that a *Dnmt1* deficiency leads to increased spontaneous firing rates of cortical neurons, reduced cortical gamma oscillations, and altered visually evoked neural responses. Despite intact sensory perception, *Dnmt1*-deficient mice exhibited reduced physical activity, heightened anxiety-like behavior, and signs of anhedonia and apathy, which represent core features of MDD. These results underscore the critical role of DNMT1 in PV interneuron function and identify it as a potential target for MDD research.

## 1 Introduction

GABAergic interneurons are crucial for healthy brain function by regulating the activity of cortical circuits. Abnormal GABAergic circuits cause changes in synaptic responses, affecting pyramidal neuron output and contributing to disease-related behaviors^1–3^. Dysfunctions of cortical inhibition are often linked to neuropsychiatric disorders such as schizophrenia and major depressive disorder (MDD)^4–7^. However, there are contradictory findings on GABAergic transmission and signaling in MDD, complicating a mechanistic understanding^8,9^. Moreover, neuropsychiatric diseases are multifactorial in their origin as they arise from a combination of genetic and environmental factors. The broad spectrum of symptoms further hampers attempts at pinpointing the underlying pathophysiology. Epigenetic modifications, including DNA methylation by DNA methyltransferases (DNMTs), link external factors to disease onset and progression. Altered DNA methylation profiles have been observed across various neuronal subtypes in conditions, such as schizophrenia, epilepsy, and MDD^10–14^. Specifically, altered expression of the DNA methyltransferase 1 (DNMT1) in cortical GABAergic interneurons has been linked to schizophrenia, highlighting its role in interneuron function and relevance to neuropsychiatric diseases (reviewed in^4^).

Parvalbumin-expressing (PV) interneurons represent the largest group of GABAergic inhibitory interneurons in the cerebral cortex and play a key role in maintaining the excitation/inhibition (E/I) balance^15^. Changes in the number and/or function, specifically of PV interneurons, were shown to be causal for the aforementioned neuropsychiatric diseases^16–19^. PV interneuron function is affected by chronic stress and implicated in MDD pathology^20^, but the molecular changes underlying altered PV interneuron functionality remain unclear. Our previous study demonstrated that DNMT1-dependent DNA methylation modulates synaptic transmission of inhibitory cortical PV interneurons by regulating endocytosis-dependent GABA reuptake at the presynaptic terminal^21^. Thus, DNMT1 may contribute to flexible changes in PV interneuron function, suggesting dysregulated *Dnmt1* expression in these cells to be a potential cause for disease features evident in MDD.

Building on these findings, we investigated the functional repercussions of dysregulated PV interneuron function due to DNMT1 deficiency at cellular, network, and behavioral levels. We found fundamental changes in neuronal activity on the single-cell level with clear implications for network activity. Furthermore, we observed behavioral deficits in *Dnmt1*-deficient mice indicating a dysphoric state. In sum, the electrophysiological and behavioral aberrations are reminiscent of an MDD phenotype.

## 2 Results

### 2.1 *Dnmt1*-deletion in PV interneurons alters local network activity in the primary visual cortex (V1)

DNMT1 impacts the GABAergic transmission of *Pvalb*-expressing (PV) cortical interneurons by acting on vesicle replenishment through the DNA methylation-dependent regulation of endocytosis-associated gene expression^21^. As cortical GABAergic interneuron dysfunction and dysregulated *Dnmt* expression have both been implicated in neuropsychiatric diseases^4,22–24^, we here aimed to reveal the functional consequences of *Dnmt1* deletion in parvalbumin-positive (PV) interneurons on the network level.

Pensold et al.^21^ found increased inhibitory postsynaptic currents in excitatory cortical neurons of *Pvalb-Cre/Dnmt1*-KO mice as well as elevated levels of extracellular GABA within the cortex. To directly test if this relates to changes in the spontaneous activity of cortical PV interneurons in *Dnmt1*-KO mice, we performed in vivo electrophysiological recordings using Neuropixels probes. Here, we recorded cortical activity in awake, behaving mice with or without endogenous *Dnmt1*-expression in PV interneurons (Fig. 1a; control mouse line: *Pvalb-Cre/tdTomato;* knockout mouse line: *Pvalb-Cre/tdTomato/Dnmt1 loxP^2^*). To selectively identify PV interneuron activity, a viral construct (AAV1.shortCAG.dlox.hChR2(H134R).WPRE.hGHp) was injected into the primary visual cortex (V1) to induce Cre-dependent expression of the blue light-sensitive opsin Channelrhodopsin-2 (ChR2). As the expression of Cre was restricted to PV interneurons in our mouse model, these cells could be selectively activated by blue light. This allowed us to isolate their neural activity patterns from the recorded populations of neurons based on their short-latency responses to 10-ms-long blue light pulses (Fig. 1b, action potential waveform examples). Spontaneous firing rates of PV interneurons in *Pvalb-Cre/Dnmt1*-KO mice were elevated relative to controls (Fig. 1b; *n*_Ctrl_=54 and *n*_KO_=37 neurons; Wilcoxon *rank-sum* test: *p*=0.0297), confirming that DNMT1 is crucial for regulating PV interneuron activity.

**Figure 1:**
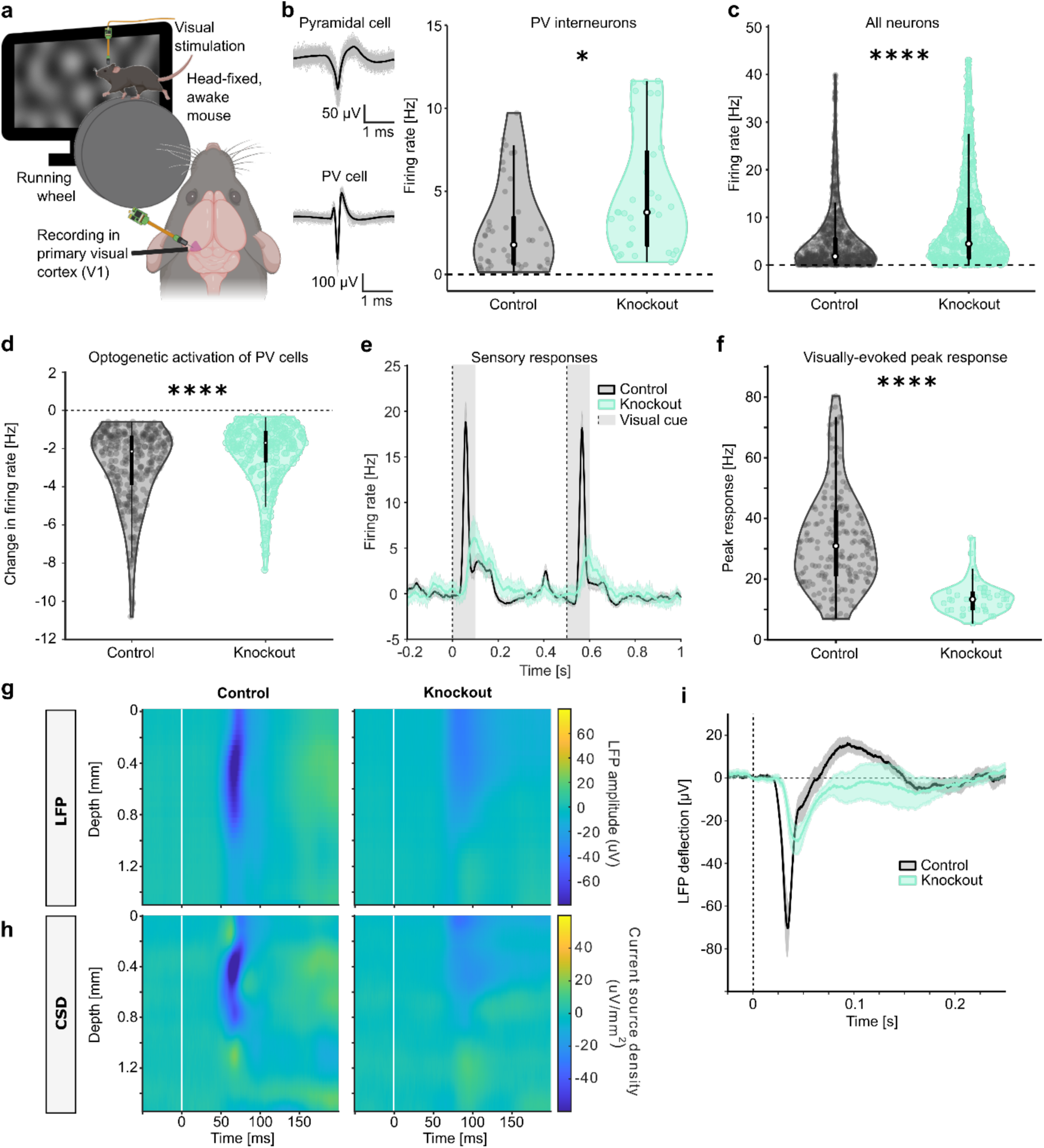
*Dnmt1*-knockout in PV interneurons alters cortical neural responses and information processing. (**a-i**) Electrophysiological recordings in the primary visual cortex (V1) using Neuropixels probes, revealed strongly altered cortical signals (*n* = 8 recordings from 2 mice per genotype). (**a**) Schematic illustration of the experimental setup and recording site. (**b**) Fast-spiking PV interneurons were identified via optotagging (see example spike waveforms, left). Their spontaneous firing rates were increased in *Pvalb-Cre/tdTomato/Dnmt1 loxP^2^* (knockout) mice. (**c**) Spontaneous firing rates of V1 neurons were significantly increased in knockout mice compared to *Pvalb-Cre/tdTomato* (control) mice. (**d**) Relative change in the firing rate of V1 neurons inhibited by optogenetic activation of PV interneurons. PV-induced inhibition was significantly weaker in knockout mice. (**e**) Average firing rates of V1 neurons in response to visual stimulation in control and knockout mice. Visual stimuli were 0.1-s-long, full-field presentations of low-frequency noise patterns, presented at time point 0 and after 0.5 seconds. Shading shows the standard error of means (SEM) across neurons. (**f**) Peak response firing rate of V1 neurons after visual stimulation. Knockout mice had significantly weaker sensory responses compared to control mice. (**g-h**) Local field potentials (LFP; **g**) and current source densities (CSD; **h**) of control and *Dnmt1*-knockout mice across the whole depth of the cortex. White lines indicate visual stimulus onset. (**i**) LFPs in V1 in response to visual stimulation at time point 0, averaged across cortical layers. Shading shows SEM across recordings. * *p* < 0.5, ** *p* < 0.01, *** *p* < 0.001, **** *p* < 0.0001.

Interestingly, when analyzing the firing rates of all cortical neurons, including all excitatory and inhibitory neurons, we also found increased spontaneous firing rates in *Pvalb-Cre/Dnmt1*-KO mice compared to control animals (Fig. 1c; *n*_Ctrl_=838 and *n*_KO_=448 neurons; Wilcoxon *rank-sum* test: *p*=5.72x10^-19^). PV interneurons inhibit excitatory neurons^25,26^, which make up the majority of cortical neurons^25,27^. The observed increase in spontaneous firing rates of all cortical neurons (Fig. 1c) could therefore be due to a homeostatic adaptation to continuously elevated inhibition caused by *Dnmt1*-deficiency in PV interneurons^21^, which could culminate in a decreased inhibitory impact of PV interneurons on cortical network activity. To test this, we exploited the optogenetic approach to activate ChR2-expressing PV interneurons, which is commonly used to inhibit brain regions in a targeted manner^28,29^. In line with this, neuronal firing rates in *Pvalb-Cre/Dnmt1*-control mice were strongly diminished upon PV cell stimulation (Fig. 1d). In contrast, inhibition in *Pvalb-Cre/Dnmt1*-KO mice using the same optogenetic PV cell activation was substantially reduced (Fig. 1d; *n*_Ctrl_=332 and *n*_KO_=238 neurons; Wilcoxon *rank-sum* test: *p*=3x10^-6^). Previous research found the number and distribution of cortical PV cells to be unaffected by the *Dnmt1*-knockout^30^. Hence, the diminished inhibitory power observed after PV activation in *Pvalb-Cre/Dnmt1*-KO mice compared to control mice likely represents a functional adaptation of the network in response to altered synaptic efficacy of PV neurons. Such a reduced impact of PV interneurons onto their postsynaptic target cells would also explain the elevated spontaneous firing rates determined for all cortical neurons (Fig. 1c).

To evaluate the consequence of this altered circuit function for cortical information processing, we recorded from the same neurons while presenting visual stimuli to the mice (Fig. 1a, e-i). Overall, we observed an altered network response to visual stimulation, comprising an attenuated response amplitude, reduced temporal precision, and a vastly diminished inhibitory component of cortical network responses in *Pvalb-Cre/Dnmt1*-KO mice compared to controls (Fig. 1e-i, Supp. Fig. 1a-c). Specifically, using 100-ms-long presentations of full-field low-frequency visual noise stimuli (see Fig. 1a), we reliably elicited strong cortical responses in V1 of control mice (Fig. 1e). In contrast, cortical responses were significantly reduced in *Pvalb-Cre/Dnmt1*-KO mice (Fig. 1e-f; *n*_Ctrl_=293 and *n*_KO_=74 neurons; peak responses in panel f were compared using Wilcoxon *rank-sum* test: *p*=1.1x10^-15^). Cortical responses in *Pvalb-Cre/Dnmt1*-KO mice to visual stimulation also arose with a temporal delay, reaching the response peak more than 50 ms later than control neurons (Supp. Fig. 1a; Wilcoxon *rank-sum* test: *p*=2x10^-26^). This loss in temporal precision of sensory responses was observed across all visually responsive neurons, with the full width at half maximum being more than twice as broad in *Pvalb-Cre/Dnmt1*-KO mice compared to controls (Supp. Fig. 1b; Wilcoxon *rank-sum* test: *p*=1.4x10^-7^). We also calculated the area under the receiver operating characteristic (ROC) curve (AUC) to quantify how distinct the change in neural activity during visual stimulation was compared to baseline conditions (Supp. Fig. 1c). Here, values closer to 0.5 indicate no differences in activity during stimulation while values closer to 1 represent perfect separation of stimulus-driven and baseline activity. In line with the weaker sensory responses in *Pvalb-Cre/Dnmt1*-KO mice (Fig. 1e), we found fewer stimulus-responsive neurons (*n*_Ctrl_=31%, *n*_KO_=22%; Wilcoxon *rank-sum* test: *p*=6.1x10^-4^), responding with a lower specificity (Supp. Fig. 1c; Wilcoxon *rank-sum* test: *p*=1.4x10^-4^). Local field potentials (LFPs; Fig 1g) and current source densities (CSD; Fig. 1h) for control and *Dnmt1*-KO mice also showed that the reduced excitatory and inhibitory responses in *Pvalb-Cre/Dnmt1*-KO mice were found across all cortical layers. In line with earlier findings^31^, visual responses in control mice showed a clear spatial structure, with the earliest and most pronounced responses in layer 4 (0.3 – 0.5 mm) and subsequent activation of the superficial and infragranular layers. In contrast, sensory responses in *Pvalb-Cre/Dnmt1*-KO mice were much weaker and less spatially defined, suggesting a disruption of layer-specific information processing. In line with the role of PV interneuron activity in decisively shaping cortical network responses and the impaired PV interneuron activity upon *Dnmt1* deletion (Fig. 1e), the neural response in *Pvalb-Cre/Dnmt1*-KO mice was strongly altered across all layers and did not exhibit an inhibitory overshoot component that would be indicative of feedback inhibition (Fig. 1i). In sum, these results imply that *Dnmt1* deletion in PV interneurons strongly reduces temporal precision and weighting of the cortical response towards visual stimulation in awake mice.

### 2.2 Dissociation between impaired cortical neural function and corresponding behavioral outcomes

Next, we aimed to test, whether impaired visual responses would translate into measurable changes in visual perception. Exploiting the mice’s optomotor response to moving gratings of differing spatial frequencies and contrasts, we first tested their visual acuity and contrast sensitivity (Supp. Fig. 1d-e). However, *Pvalb-Cre/Dnmt1*-KO mice displayed no impairments compared to controls (visual acuity tested with ANOVA: *F*=0.039, *df*=1, *p*=0.84; contrast sensitivity: *F*=0.0103, *df*=1, *p*=0.919). This implies that basic sensory processing remains largely intact despite the molecular and cellular effects induced by the PV-specific *Dnmt1*-knockout that trigger alterations in cortical activity. It is noteworthy that optometry performances additionally rely on subcortical structures^32^, which could partly compensate for defective cortical functionality^33^.

Thus, we next aimed to investigate whether *Dnmt1-*deficiency in PV interneurons influences behavioral outcomes of more complex visual tasks that rely on higher-order cortical dynamics. For this, we focused on the anterolateral motor cortex (ALM). The ALM is located in the prefrontal cortex (PFC) and plays a key role in integrating sensory information with motor planning and decision-making, particularly in tasks requiring visual perception, learning, and evidence accumulation^34–36^. While its primary role is movement-preparation, the ALM can acquire responsiveness to behaviorally relevant visual stimuli in a context-dependent manner^34,37^, especially in tasks requiring sensorimotor integration (e.g., when visual stimuli inform learned motor responses). To measure local network activity in response to visual stimulation in the ALM, we conducted Neuropixels recordings, in mice that were trained to perform a licking response (motoric reaction) toward the same side where a visual stimulus was presented. Visual stimuli were moving sinusoidal gratings on one of two lateral monitors to either the right or the left side of the animal (Fig. 2a, scheme). Mice of both genotypes equally learned the task, demonstrating that *Dnmt1* deletion in PV interneurons did not induce clear learning deficits (Supp. Fig. 2a).

**Figure 2:**
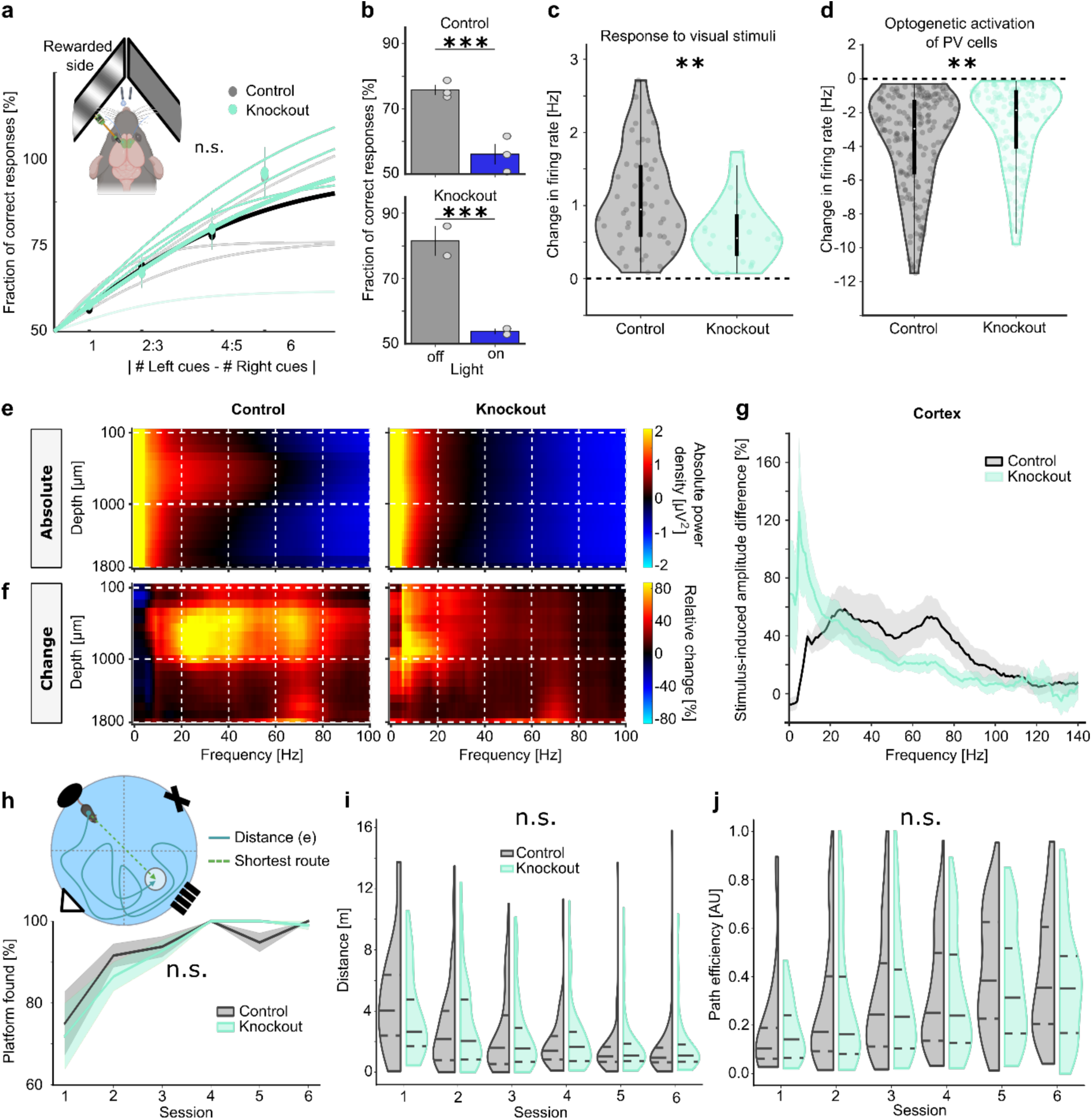
*Pvalb-Cre/tdTomato/Dnmt1 loxP^2^* (knockout) mice display intact behavioral outputs despite their changes in gamma oscillations and prefrontal cortex (PFC) activity. (**a-d**) Electrophysiological recordings in the anterolateral motor cortex (ALM) of mice trained on a visual evidence accumulation task showed similar changes in cortical activity as in V1 (*n*_(Control)_ = 3, *n*_(Knockout)_ = 4). (**a**) Mice were trained to detect visual stimulation on one of two screens to later perform an evidence accumulation task and record activity in ALM (scheme on the upper left side). In this visual evidence accumulation task, mice (*n*_(Control)_ = 3, *n*_(Knockout)_ = 4) reported the side with the higher number of cues by licking the corresponding water spout. Knockout mice displayed no behavioral differences regarding their performance in this task (graph). (**b**) Optogenetic activation of PV cells in the ALM likewise impaired decision-making in control (*n* = 3, top) and knockout (*n* = 2, bottom) mice. (**c**) Firing rates of ALM neurons in response to visual stimulation in trained mice showed weaker signals in knockout compared to control mice. (**d**) Baseline-corrected firing rate of all PFC neurons inhibited during optogenetic activation of PV interneurons. PV-induced inhibition was significantly weaker in knockout mice compared to control animals. (**e-g**) Knockout mice displayed strongly diminished gamma oscillations (30–90 Hz) in comparison to *Pvalb-Cre/tdTomato* (control) mice (*n* = 8 recordings from 2 mice per genotype). (**e**) Absolute spectral power density during visual stimulation with full field moving gratings. Visual stimulation elicited high-frequency oscillations in the primary visual cortex (V1) of control but not knockout mice. Colors show spectral power on a log_10_ scale. (**f**) Relative change in spectral power compared to baseline activity. Visually evoked oscillations in control mice matched the gamma band (30–90 Hz), while in knockout mice oscillations of much lower frequencies (< 20 Hz) were measured. (**g**) Change in spectral power within the cortex displayed as mean ± SEM. The direct comparison between control and *Dnmt1*-knockout mice revealed the shift towards lower-frequency oscillations in knockout mice. (**h-j**) Learning and memory formation were tested in a Morris water maze (MWM), showing similar performances of both genotypes (*n* = 8 mice per genotype). (**h**) Mice were introduced into the MWM from randomly alternating sites and had to find a hidden platform based on visual landmarks (upper scheme). Mice of both genotypes similarly improved their success rate across sessions (lower graph, mean ± SEM of mice). Both the swimming distances (**i**) and the efficiencies of the chosen paths (distance/shortest route, **j**) did not differ in knockout mice. * *p* < 0.5, ** *p* < 0.01, *** *p* < 0.001, **** *p* < 0.0001.

In agreement with earlier studies^35,36^, optogenetic activation of PV interneurons in ALM using viral injections of AAV1.shortCAG.dlox.hChR2(H134R).WPRE.hGHp and blue light stimulation also strongly reduced the choice accuracy of trained control mice, demonstrating that the ALM was required for solving the task (Fig. 2b, top). The same choice impairment was also found in *Dnmt1*-knockout mice, suggesting that the ALM retained its importance for decision-making in these animals (Fig. 2b, bottom). Indeed, Neuropixels recordings in task-performing animals of both groups revealed neuronal responses to visual stimulation in the ALM, confirming successful visuomotor learning (Fig. 2c). Similar to our observations in V1, sensory responses in *Pvalb-Cre/Dnmt1*-KO mice were significantly reduced compared to control mice (Fig. 2c; *n*_Ctrl_=49 and *n*_KO_=29 neurons; Wilcoxon *rank-sum* test: *p*=2.9x10^-3^). This difference might either be a consequence of the impaired processing in visual cortex or suggests that cortical network alterations due to disrupted PV interneuron function also generalize to the frontal cortex. To directly test this, we again used blue light stimulation to optogenetically trigger PV activity in the ALM. Indeed, the neural firing of ALM neurons was significantly less affected by the optogenetic stimulation of PV neurons in KO versus control mice (Fig. 2d; *n*_Ctrl_=196 and *n*_KO_=158 neurons; Wilcoxon rank-sum test: *p*=2.5x10^-5^), despite the fact that the PV cell distribution in the frontal cortex of *Dnmt1*-KO versus control mice was largely similar (Supp. Fig. 3). This is in line with a reduction of the efficacy of cortical inhibition by PV interneurons, suggesting that the same microcircuit changes found in V1 also apply to higher-order cortices.

To test whether the observed higher-order functional changes at the cellular level translate into related behavior, we conducted a visual evidence-accumulation task. Here, head-fixed animals were presented with visual cues on both of the two screens, and had to indicate on which screen a higher number of visual cues was presented throughout a 3-s stimulation period, by licking the corresponding waterspout after a 0.5-s delay. Such accumulation of sensory information was shown to rely on the interplay between sensory and frontal cortical areas^38–40^. However, performances in this visual evidence-accumulation task were not affected by the *Dnmt1*-deletion in PV interneurons (Fig. 2a; LME model: *p*=0.85, *n*=30 performance values from 10 mice over 3 difficulties).

The observation that *Pvalb-Cre/Dnmt1-*KO mice do not display behavioral deficits despite significant electrophysiological changes, suggests the presence of compensatory mechanisms that keep sensory perception intact.

### 2.3 *Dnmt1*-KO in PV cells induces changes in gamma oscillations without affecting learning and memory

PV interneuron activity is known to be involved in the generation of cortical gamma oscillations^19,41,42^. Interestingly, we found that the *Dnmt1*-KO resulted in severely reduced visually evoked gamma oscillations (Fig. 2e-g) in addition to the altered firing patterns in V1. Gamma oscillations are required for the synchronization of different cortical areas to enable complex processes such as learning and memory formation^43^. Therefore, we next sought to investigate potential deficits in learning and memory capacities of the *Dnmt1*-KO mice using a common learning task, the Morris water maze (Fig. 2h-j). We placed mice into a pool of clear water from alternating quadrants and quantified their ability to repetitively locate a submerged, translucent platform based on several visual cues on the arena walls. As optometry pointed to an intact visual perception, we decided on this memory task involving visual navigation. Overall, we observed no significant differences in the performance of *Pvalb-Cre/Dnmt1*-KO mice compared to controls in the Morris water maze (Fig. 2h-j). *Pvalb-Cre/Dnmt1*-KO mice were able to locate the platform as often (Fig. 2h; Binomial test: *p*=0.716) and similarly fast and effective as control animals (Fig. 2i-j; Distance: Two-Way ANOVA (Repeated Measures): *p*(Genotype)=0.63, *p*(Sessions)=1.3x10^-22^, *p*(Genotype:Sessions)=0.92; Path efficiency: Two-Way ANOVA (Repeated Measures): *p*(Genotype)=0.11, *p*(Sessions)=7.9x10^-15^, *p*(Genotype:Sessions)=0.36). In line with these data, learning curves in the visual evidence accumulation task described above were also similar for *Dnmt1*-KO and control mice (Supp. Fig. 2a). This result was further reinforced by the learning curves of *Pvalb-Cre/Dnmt1*-KO and control mice obtained in a touchscreen chamber setup, where freely moving mice had to learn to touch the screen with a visual stimulus (Supp. Fig. 2b). Moreover, in line with the optometry results, the touchscreen chamber test further confirmed intact visual perception. Here, a cohort of animals was trained to discriminate a black circle from a lower-contrast distractor stimulus by touching the respective side of the screen (Supp. Fig. 2c, upper scheme). This test likewise revealed no significant differences between *Pvalb-Cre/Dnmt1*-KO and control mice (Supp. Fig. 2c, graph; ANOVA: *F*=0.45, *df*=1, *p*=0.501). All mice achieved comparable expert performances for different difficulty levels, i.e., different contrast levels of the distractor stimuli.

In sum, similar learning capacities of mice with and without endogenous *Dnmt1*-expression in PV interneurons were observed in different experimental setups (Fig. 2h-j, Supp. Fig. 2). Thus, the change in gamma synchrony did not affect the animals’ abilities to learn and memorize different sensory integration-based tasks.

### 2.4 *Dnmt1*-deficient mice display reduced physical activity and increased body weight

So far, electrophysiological changes have neither translated into impaired sensory perception, learning deficits or defective task-solving abilities involving higher-order brain structures. Of note, impaired gamma oscillations were observed in patients with mood disorders, such as depression or bipolar disorder, and constitute a hallmark in mouse models of depression^19,44^. Moreover, the ALM displayed electrophysiological changes in *Dnmt1* deficient mice. The ALM is located in the PFC, and GABAergic deficits and dysfunctions in this specific region have been implicated in symptoms related to neuropsychiatric diseases such as depression^20^ and anxiety-related behavior^1,2,18,20^. Different methods altering the activity of PFC circuitry and projections from PFC to other brain regions changed the behavioral output of mice in depression-related tasks. Interestingly, RNA-sequencing results of *Pvalb-Cre/Dnmt1*-KO mice also indicated gene expression changes matching those found in different mouse models expressing behavioral traits of depression and anxiety disorder (Supp. Tab. 1-2). This led us to investigate whether *Dnmt1*-deficiency in PV interneurons is associated with signs of mood disorders.

Analyzing the activity of *Pvalb-Cre/Dnmt1*-KO and control mice in their home cage, an undisturbed and familiar setting, unveiled a consistent reduction in the overall movement of *Pvalb-Cre/Dnmt1-*KO animals (Fig. 3a-b). For this, we monitored naive animals in their home cages and chose several uninterrupted 30-min time intervals per day to compare the animals’ activity during these periods. By calculating the area of differing pixels between every fifth frame of the video, we obtained a measure of their movement (Fig. 3a). Across all observed intervals, *Pvalb-Cre/Dnmt1-*KO mice consistently displayed lower levels of activity (Fig. 3b; two-way ANOVA: *p*(Genotype)=1.9x10^-5^, *p*(Time Interval)=0.98, *p*(Genotype:Time Interval)=0.91). Causes for such a change in unguided behavior can be both physiological (e.g., due to muscle impairments) or psychological. In the latter case, physical movement could be altered as a result of reduced intrinsic motivation or fatigue due to mood disorders such as depression. Exploration, movement, and interaction with enrichments are regular behaviors found in mice and a reduction of these activities can arise from various factors. Therefore, we conducted a battery of tests to rule out physical causes. Based on the results described above, visual impairments can be excluded as an underlying cause for decreased activity (see Fig. 2a, Supp. Fig. 1d-e, 2). Alternatively, reduced physical activity could result from impairments of the motor system. In a previous study, we found that muscular innervation as well as muscle integrity and structure in *Pvalb-Cre/Dnmt1*-KO mice are intact^30^. Similarly, somatomotoric performance, tested in a ladder rung test, revealed no defects due to *Dnmt1* deletion in *Pvalb-Cre*-expressing cells in the relevant age groups^30^. We additionally performed a rotarod test and a wire hang test to further screen for potential defects in locomotor activity at the behavioral level. While no significant differences between knockout and control animals were observed in the latency to fall during a rotarod test (Fig. 3c; ANOVA: *F*=1.037, *df*=1, *p*=0.31), we found a reduced latency to fall for *Dnmt1*-knockout mice in the wire hang test (Fig. 3d; ANOVA: *F*=39.62, *df*=1, *p*=9.8x10^-9^). Notably, the wire hang test examines general motor function but depends on various factors such as age, sex, and body weight. Hence, we measured and compared body weights of DNMT1-deficient and control mice. Strikingly, the body weight of *Pvalb-Cre/Dnmt1-*KO mice was significantly increased (Fig. 3e; Student’s *t*-test: *p*=1.8x10^-6^), which could explain the strongly reduced latency to fall in the wire hang test, despite all other muscle and motor analyses being inconspicuous.

**Figure 3:**
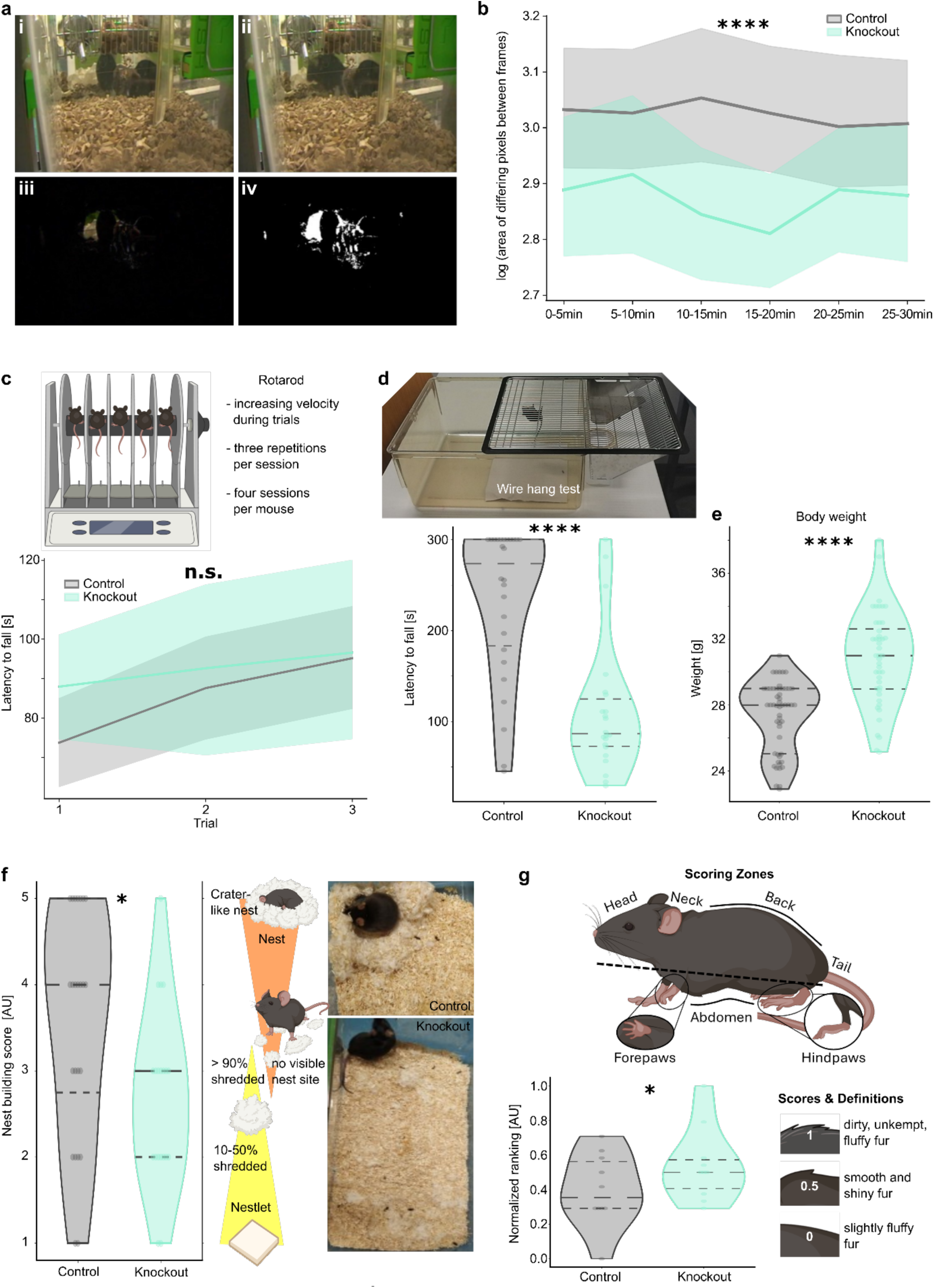
*Pvalb-Cre/tdTomato/Dnmt1 loxP^2^* (knockout) mice display apathetic behavior. (**a-b**) Spontaneous activity within the home cage was consistently diminished in knockout mice compared to *Pvalb-Cre/tdTomato* (control) mice (*n* = 6 mice per genotype). (**a**) Mice were monitored in their home cages to evaluate their spontaneous, undisturbed behavior. By comparing every fifth frame from the resulting videos (i, ii), the change in pixel values between frames (iii) was binarized (iv) to calculate the change in position of the contained animals. (**b**) A comparison across multiple time intervals showed an overall reduced average activity of knockout mice compared to controls (shading indicates SEM). (**c**) Average latency to fall from a rotating rod (‘Rotarod’) with increasing velocity showed no apparent motor deficits in knockout mice (*n*_(Control)_ = 13, *n*_(Knockout)_ = 11). Results from three consecutively conducted trials are shown with shadings indicating the SEM. (**d**) A wire hang test (top image) showed a significantly reduced latency to fall in knockout mice (bottom graph) which can indicate a difference in motor strength but is highly dependent on the animals’ body weight (*n*_(Control)_ = 13, *n*_(Knockout)_ = 11). (**e**) Increased body weights were elevated for knockout compared to control mice (*n*_(Control)_ = 13, *n*_(Knockout)_ = 11). (**f**) Nest-building capacities were significantly decreased in KO (*n* = 25) versus control mice (*n* = 24). As the representative images (right side) show, knockout mice were shredding the provided material but failed to assemble visible nest sites. (**g**) Assessment of the animals’ coat state as indicated in the illustration on the bottom right showed significantly higher rankings of the knockout mice, indicating worsened fur care (*n* = 12 mice per genotype). * *p* < 0.5, ** *p* < 0.01, *** *p* < 0.001, **** *p* < 0.0001.

As there is a relation between increased body weight and mental health, i.e., a comorbidity of weight gain and other depressive symptoms^45^, we further investigated the cause for the observed weight differences in more detail. The weight differences could either result from the reduced activity of *Pvalb-Cre/Dnmt1*-KO mice, leading to diminished energy expenditure, or from altered eating behavior. Therefore, we next examined whether *Dnmt1*-KO mice showed indications of altered food intake or changes in appetite regulation.

We first checked the animals’ food consumption in their home cages of grouped and single-housed mice by assessing the change in pellet weight normalized to the number of mice after 24 h and 72 h. For both experimental groups, single- and group-housed mice, no difference in food intake of *Dnmt1*-knockout mice compared to control animals was observed (Supp. Fig. 4a-b). On the contrary, food and water intake in relation to the respective animals’ body weight were even significantly decreased for *Pvalb-Cre/Dnmt1*-KO mice (Supp. Fig. 4c). These results render the food and water intake unlikely parameters to explain the difference in weight gain. However, these experiments provide only a brief glimpse at the animals’ consummatory behavior. Therefore, we proceeded by investigating hormonal concentrations (Supp. Fig. 4d) and the subcortical architecture of select areas (Supp. Fig. 5-6) to compare markers of appetite regulation.

Body weight control essentially depends on hormones circulating in the blood and its regulatory neuronal pathways as well as peripheral mechanisms that not only control food intake, but also energy storage and expenditure^46,47^. Appetite-regulating hormones can either act orexigenic (promoting appetite) or anorexigenic (reducing appetite). These hormones are detected by neurons of the hypothalamus, in particular the arcuate nucleus^46,48^, which is interconnected with diverse brain areas. Together, they orchestrate appetite and feeding control with implications for body weight regulation^49,50^. Thus, we measured the blood concentrations of appetite-relevant hormones in the blood of *Pvalb-Cre/Dnmt1*-KO and -control mice (Supp. Fig. 4d). Apart from leptin, which is described to act anorexigenic^46^, we did not find altered levels of hormones that regulate food intake. The orexigenic hormone ghrelin, which is produced and released from the gut and known to increase food intake by binding to receptors in the hypothalamus^51^, was not altered (Supp. Fig. 4d). The blood glucose level-regulating hormones insulin and glucagon were likewise unchanged (Supp. Fig. 4d). Both insulin and leptin work closely together to activate anorexigenic neuronal pathways^46^. In addition, the levels of glucagon-like-peptide-1 (GLP-1), which is secreted after food intake and stimulates the release of insulin from the pancreas while suppressing the secretion of glucagon, were likewise not different in *Pvalb-Cre/Dnmt1*-KO compared to control mice. Peptide YY (PYY) levels, which is co-secreted with GLP-1 and described to have an anorexigenic effect^52^, was also not altered in conditional *Dnmt1*-knockout mice (Supp. Fig. 4d).

Of note, leptin is mainly produced in adipose tissue and its levels in the bloodstream correlate with the amount of body mass^53,54^. Thus, the increased leptin levels in *Pvalb-Cre/Dnmt1*-KO mice likely represent a mere consequence of the increased body weight rather than being its cause, since leptin activates anorexigenic neuronal pathways^46^.

In sum, our findings did not point to altered food intake or hormonal states that act on food intake in *Pvalb-Cre/Dnmt1*-KO mice. These findings are in line with the very low numbers of *Pvalb-*positive cells present in the arcuate nucleus in both *Dnmt1*-KO and control mice (Supp. Fig. 5). In the arcuate nucleus, two cell groups have been described extensively: AgRP (agouti-related protein)- and POMC (proopiomelanocortin)-expressing neurons. While AgRP neurons are the main mediator of orexigenic pathways driving increases in food consumption, POMC neurons work in the opposite direction^50^. Of note, subpopulations of POMC-expressing neurons not only influence feeding but also other functions that might influence body weight, such as physical activity^55,56^.

Therefore, we next determined whether PV cells in the arcuate nucleus belong to the POMC and/or AgRP cell population, using a β-endorphin antibody for immunostaining. β-Endorphin is a product of POMC and has been described to work well for marking POMC-expressing neurons in the hypothalamus^57,58^. While a cellular signal was detected in the more anterior parts of the arcuate nucleus (Supp. Fig. 6a), as well as in synapses, and axonal fibers throughout the whole arcuate nucleus, no overlap was detected with the *tdTomato*-positive cells, neither in the soma nor in the fibers. This result indicates that PV is not expressed in POMC-expressing neurons, thus *Dnmt1* expression in these neurons should not be affected.

To control whether PV cells belong to the AgRP-expressing cell population, which often express GABA^59^, a GABA immunolabeling was performed. We found a co-localization with GABA in some but not in all tdTomato-expressing cells (Supp. Fig. 6b). While AgRP-cells promote appetite, their activity is inhibited by leptin. The low number of *Pvalb-Cre/tdTomato*-positive cells in the arcuate nucleus therefore seems to be an unlikely explanation for lasting changes in body weight regulation.

In sum, these results suggest that the increased body weight of *Pvalb-Cre/Dnmt1*-KO mice is not due to impaired orexigenic or anorexigenic hypothalamic pathways, as the PV cell distribution was similar in both genotypes and their identity was insufficiently determined, with only some PV cells potentially belonging to the AgRP neuron population. Given that single-cell sequencing approaches of the energy balance-relevant hypothalamic arcuate–median eminence complex did not indicate significant *Pvalb*-expression levels in any identified cell group^60^, the *Dnmt1*-KO in our mouse model is likely to bear no consequences for cells involved in appetite regulation. Furthermore, nutritionally relevant hormone levels showed no genotype-dependent aberrations. In support of this, no differences were seen in absolute food consumption, suggesting that hypothalamus-mediated appetite and food intake controls were not affected by *Dnmt1*-deletion in PV cells. Together, this suggests that rather the reduced energy expenditure due to diminished motor activity caused the observed weight gain in *Pvalb-Cre/Dnmt1*-KO.

### 2.5 *Dnmt1*-deficient mice display apathy- and anhedonia-like behavior

Reduced motor activity and increased body weight point to lower innate motivation as seen in an apathy-like state^61^. A common test for apathetic behavior in rodents is to assess their nest-building abilities^62^. Nest-building represents an innate behavior of mice which is required for heat conservation and shelter. Depending on environmental influences, drug intervention, or genetic manipulation, nest-building quality can deteriorate, underlining this test’s sensitivity to the animals’ health and welfare. Using an ordinal scaling system from 1-5, we found nest scores of *Dnmt1*-knockout mice to be significantly lower (Fig. 3f; Mann-Whitney *U* test: *p*=0.034). In fact, the mean and median nest scores suggest that *Dnmt1*-deficient mice shredded the provided nesting material but did not build a nest site within a 16-hour time window, suggesting lower innate motivation.

Assessing the coat state of mice provides further information about their overall health and well-being^63,64^ and deterioration of coat state also points to apathetic mouse behavior^65^. Therefore, we conducted a detailed analysis, assessing the fur state of seven different body regions of the same individuals in a blinded manner as described by Nollet et al.^64^. This revealed a systematic neglect of the coat state of *Pvalb-Cre/Dnmt1*-KO mice (Fig. 3g). While alterations in coat state were mostly moderate in both genotypes, such as slightly matted or spiky patches of fur in isolated body areas, *Pvalb-Cre/Dnmt1*-KO mice displayed these alterations in more body regions. Thus, their cumulative score was significantly increased, indicating a deterioration of their overall coat care in comparison to control mice (Fig. 3g).

With the phenotype characterization so far pointing towards decreased levels of innate motivation, we next investigated the animals’ engagement in rewarding behaviors. First, we analyzed voluntary wheel running of *Pvalb-Cre/Dnmt1*-KO and -control mice in their home cages. We quantified running activity in individual mice for up to three consecutive days after animals were separated from cage mates for at least one week. In line with the reduced movement in home cages described above, we observed that *Dnmt1*-KO mice displayed a significant reduction in running activity (Fig 4a-c). As expected, activity levels of all tested animals largely varied depending on the light cycle, with much more frequent wheel running during night times (Fig. 4b). Regardless of the time of day, *Dnmt1*-KO mice ran significantly less than control animals (Fig. 4b), with differences being particularly pronounced during the dark phase (Fig. 4c; Wilcoxon *rank-sum* test: *p*=0.843x10^-14^), suggesting a greatly reduced interest in the provided wheel. As changes in voluntary wheel running activity indicate alterations in motivation, mood, and overall well-being^66^, the diminished usage of the running wheel could reflect reduced motivation for rewarding behavior^67^, which is a hallmark feature of depression^68^.

**Figure 4:**
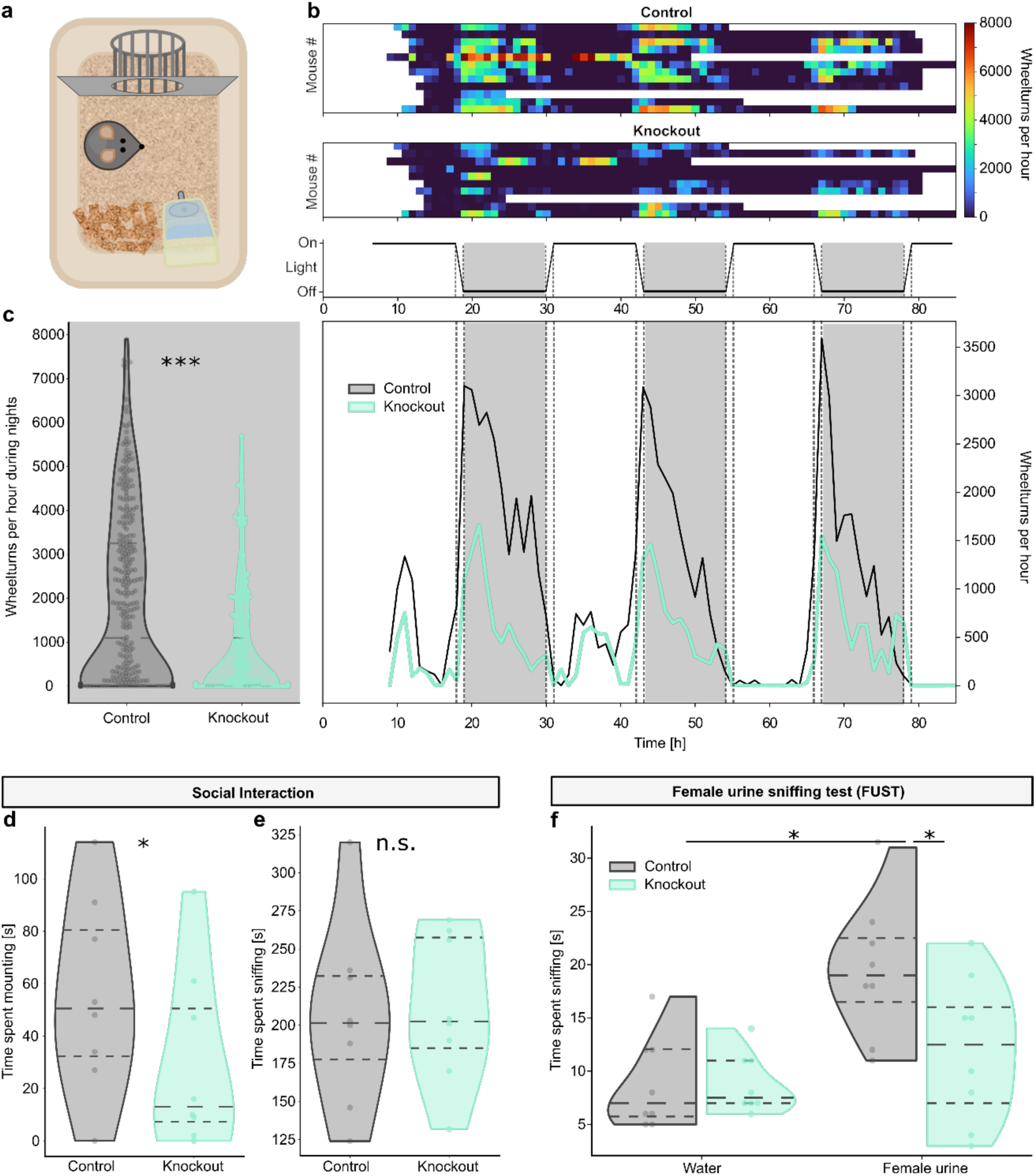
*Pvalb-Cre/tdTomato/Dnmt1 loxP^2^* (knockout) mice show reduced self-motivated behavior, indicating an anhedonic state. (**a-c**) Voluntary running on a wheel provided within the home cage was strongly reduced in knockout (*n* = 9) versus *Pvalb-Cre/tdTomato* (control) mice (*n* = 11). (**a**) Schematic illustration of the voluntary running experiment. Running wheels were provided in the home cages of single-housed mice for up to three nights. (**b**) Almost all mice displayed fluctuating utilization of the running wheel (upper heat maps) based on the daytime (middle graph), but in general, knockout mice were running less frequently than controls. The overall reduced activity of knockout mice is discernible in the mean running time averaged per genotype (lower graph). (**c**) Quantifying the running activity during the animals’ active period at night showed a significant reduction in running wheel usage of knockout versus control mice. (**d-e**) Evaluating the interaction of male knockout and control mice with female C57BL/6J-mice in a 10-min time frame reveals social interaction with female conspecifics (*n* = 8 mice per genotype). (**d**) knockout mice spent significantly less time mounting the females than controls. (**e**) Males of both genotypes spend an equal time interacting with a novel female conspecific. (**f**) Presentation of urine from female C57B/6J mice significantly increased time spent sniffing in control but not in knockout mice (*n* = 8 mice per genotype). * *p* < 0.5, ** *p* < 0.01, *** *p* < 0.001, **** *p* < 0.0001.

In summary, *PV-Cre/Dnmt1*-KO mice display a loss of innate motivation in various biologically relevant contexts. Such reduced participation in otherwise regularly executed activities indicates an apathy-like state and can also be accompanied by anhedonic behavior. Anhedonia, a core symptom of depression, refers to a reduced ability to experience pleasure or interest in activities that are usually considered enjoyable^69^. Considering that voluntary wheel running in mice was shown to be a modifier of the dopamine system^70^ causing an antidepressant-like effect^71^, reduced running activities could also be driven by a changed hedonic valuation.

Typical tests for anhedonia in rodents often exploit the animals’ valuation of palatable foods (e.g., sucrose consumption/preference test, sucrose self-administration, cookie test). As leptin was shown to alter the valuation of the rewarding value of sucrose, and as we found increased leptin levels in the *Dnmt1*-KO mice likely due to the increased weight (Supp. Fig. 4d), we chose to instead investigate the animals’ reward- and pleasure-seeking behavior by assessing their sexual interest.

For this, we observed and quantified behaviors of *PV-Cre/Dnmt1*-KO virgin males in response to their first encounters with female conspecifics. We consistently observed *Dnmt1*-deficient male mice spend less time trying to mount females (Fig. 4d; Student’s *t*-test: *p*=0.018). Nevertheless, their time interacting with or sniffing the respective female remained unaltered compared to control males (Fig. 4e). While this suggests that there is a reduced interest in sexual interaction in *PV-Cre/Dnmt1*-KO mice, the outcome of such social pairings can also vary depending on the interest of the females in the presented males. Therefore, we additionally conducted a female urine sniffing test. This test was specifically developed to circumvent appetite- and activity-related alterations in reward-seeking behavior^72^. Female urine constitutes a chemosensory cue for male mice, which is related to sexual behavior. Dopamine release from the nucleus accumbens was observed in male mice not only for active sexual behavior but also already during the sniffing of urine provided by females in estrus^72,73^. Therefore, providing control male mice with a sample of female urine usually arouses their interest, resulting in prolonged times sniffing the olfactory cue compared to water (Fig. 4f). However, the *Pvalb-Cre/Dnmt1*-KO mice did not display elevated interest in female urine compared to water but spent significantly less time sniffing the female urine compared to control animals (Fig. 4f; Student’s *t*-test: *p*=0.041).

Overall, we observed several behavioral patterns indicating apathy and anhedonia of *Pvalb-Cre/Dnmt1*-KO mice. Together, they indicate a dysphoric mental state of the animals as found in mood disorders, such as depression.

### 2.6 *Dnmt1*-deletion in PV interneurons leads to increased anxiety in mice

Another symptom associated with mood disorders is anxious behavior. To test for this, we conducted a novelty-suppressed feeding test (Fig. 5). Here, we examined the conflict between a desire for food versus the uncertainty of a new and exposed surrounding (Fig. 5a). For this, mice were temporarily food deprived for 24 hours. Despite the significant difference in body weight between the *Pvalb-Cre/Dnmt1-* knockout mice and the respective controls, the deprivation resulted in similar rates of body weight reduction (Supp. Fig. 7a), implying a similarly strong motivation to obtain food. Yet, *Pvalb-Cre/Dnmt1-*knockout mice displayed increased latencies to approach and consume a pellet placed in the center of an open arena (Fig. 5d; Student’s *t*-test: *p*=0.018). Furthermore, the *Dnmt1*-deficient animals’ foraging was significantly slower and closer to the walls of the arena as indicated by a generally increased distance to the pellet (Fig. 5b-c, f-g). To confirm that these behavioral differences resulted from novelty-induced anxiety, mice were removed from the arena as soon as they started to chew on the pellet (Fig. 5a). With the food deprivation still serving as an active motivator, the mice were then transferred to their respective home cages to assess their latency to approach the same pellet in a known environment. Now, *Pvalb-Cre/Dnmt1-*KO mice displayed significantly shorter latencies compared to control mice (Fig. 5e; Student’s *t*-test: *p*=0.019) while consuming the same amount of pellet within 5 min (Supp. Fig. 7b). This further substantiates that the pellet was a strong incentive for the food-deprived animals in both groups, but the novelty-induced anxiety differently shaped their behavior within the unknown arena.

**Figure 5:**
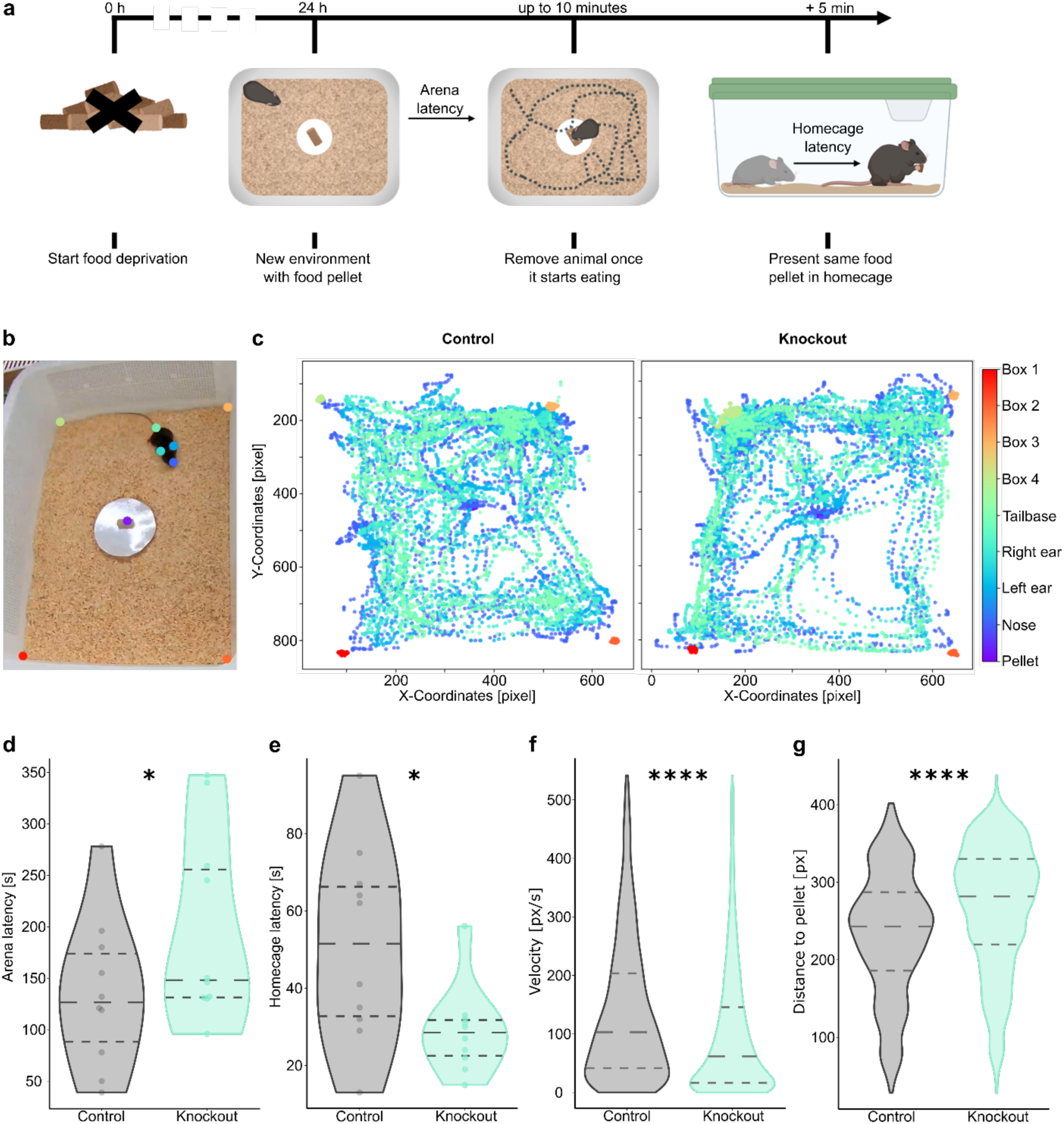
Conditional *Dnmt1* deletion in PV cells induces anxiety in a novelty-suppressed feeding (NSF) test. **a**) Scheme and schedule of the NSF test: mice (*n* = 10 per genotype) were food deprived for 24 h before being transferred into an arena with new bedding and a centrally attached food pellet of known weight. Once the animals started chewing the presented food, their latency was recorded, and they were immediately moved to their home cage. Subsequently, the pellet from the arena was presented in the home cage and the animals’ latency to approach and consume it was recorded. (**b-c**) Videos taken during the NSF test were labeled using DeepLabCut, to evaluate the animals’ behavior until they approached the food pellet. (**b**) Exemplary picture of a labeled frame. (**c**) Exemplary traces of a *Pvalb-Cre/tdTomato* (control, left) and a *Pvalb-Cre/tdTomato/Dnmt1 loxP^2^* (knockout, right) mouse show their positions within the arena during the test. (**d**) Knockout mice displayed a significantly increased latency to approach the presented food pellet in the arena compared to controls, suggesting increased levels of anxiety. (**e**) In their home cages, knockout mice approached the same food pellet significantly faster than control mice. (**f**) During their exploration of the arena, knockout mice moved with a significantly lower velocity and (**g**) a greater distance to the pellet than the controls. * *p* < 0.5, ** *p* < 0.01, *** *p* < 0.001, **** *p* < 0.0001.

Together, these behavioral results show that mice with a *Dnmt1*-knockout in PV interneurons display elevated levels of novelty-induced anxiety.

## 3 Discussion

In this study, we provide evidence that the conditional knockout of *Dnmt1* in murine PV interneurons has far-reaching consequences on brain network activity and ultimately mental health. In line with our previous results in vitro^21^, we found that a *Dnmt1* knockout in PV neurons increases their spontaneous activity, but also the overall cortical network activity in vivo, suggesting a disruption of network inhibition with implications for the cortical E/I balance. The presented data propose that a lack of network inhibition was not due to disrupted firing of PV neurons but rather a reduction in synaptic efficacy. Our earlier work showed that a *Dnmt1* deletion leads to enhanced rates of GABAergic synaptic vesicle replenishment in PV interneurons, leading to elevated GABAergic transmission^21^. The overall increase in excitation and reduction in responsiveness to PV activity that we observed here at the cortical network level therefore likely represent homeostatic adaptations of cortical pyramidal neurons that reduce their sensitivity to GABA and therefore diminish the inhibitory power of PV neurons on neural network activity. In accordance with this hypothesis, regulatory adjustments of neuronal activity to maintain E/I balance have been found in neuronal cultures but also in in vivo studies^74–76^. Similarly, the increased firing rates of PV cells could also represent such a homeostatic effect, as cortical PV interneurons have self-synapses, so-called autapses, which are crucial for the coupling of PV cell activity to induce gamma oscillations^77^. Adaptive behavior towards increased GABAergic release and elevated firing rates of neighboring neurons could both constitute modulatory mechanisms to adjust the network activity to a new status quo. Aside from an elevation of spontaneous activity, these extensive changes in cortical network properties also had a strong impact on the cortical processing of sensory stimuli, leading to a reduction in neural response amplitude and temporal precision upon visual stimulation.

Surprisingly, these clear changes in network activity were not reflected in impaired goal-directed behavior. Despite the weaker and less precise network responses to sensory stimuli, *Pvalb-Cre/Dnmt1*-KO mice performed equally well in several behavioral tasks, requiring sensory perception and discrimination. With sensory information processing also taking place in various subcortical structures^78,79^, it is conceivable that these areas compensate for imprecise cortical processing^80–82^. Similarly, learning and memory formation remained intact despite the strong distortion in cortical network activity, which could partly also be attributed to the workings of subcortical structures^83–85^. However, especially short-term memory and evidence accumulation tasks were shown to also involve neocortical function^35,36^. In compliance with the known critical role of prefrontal area ALM for short-term memory tasks^35^, we found that optogenetic inactivation of ALM similarly impeded decision-making during the visual evidence accumulation task in mice of both genotypes. Thus, it is likely that the observed changes in firing rates of the pyramidal cells and other interneurons constitute a sufficient functional adaptation to sustain overall cortical functionality despite the altered GABAergic release by PV interneurons.

It is important to note that these altered firing patterns were also present in the prefrontal cortex and manifested in reduced cortical gamma synchrony. Both of these alterations are also clinically relevant in neuropsychological diseases^18,20,24,44,86,87^. Especially PV interneurons have been noted to profoundly impact cortical gamma oscillations in the context of depression^42^. Correspondingly, we observed several behavioral abnormalities such as reduced physical activity, apathetic and anhedonic behavior, as well as increased anxiety in *Dnmt1*-KO mice. The combination of these symptoms is indicative of neuropsychiatric disorders, such as MDD or anxiety disorder^88^. Hence, while several cortex-involving processes, such as sensory signal integration, learning, and memory formation and retrieval remained functional upon *Dnmt1* deletion in PV cells, we found several behavioral abnormalities indicating depression-like symptoms.

The importance of GABAergic signaling for the pathology of neuropsychiatric diseases has been described in numerous studies^8,24,89^. Of note, the direction and scale of such changes were not necessarily congruent across studies and sometimes even seemed contradictory. Proton magnetic resonance spectroscopy revealed a general trend towards decreased in vivo GABA levels in brains of patients diagnosed with autism spectrum disorder or MDD, while no significant changes were reported for other neuropsychiatric disorders like schizophrenia or bipolar disorder^90^. The connection of decreased GABA levels to the clinical picture of MDD is further substantiated by the finding that no differences between remitted MDD patients and healthy individuals were found, suggesting a direct connection of GABA levels to symptom expression^90^. While GABA is the common transmitter of various neuron types, further assigning subtype-specific effects on disease progression often proved to be difficult. In postmortem studies, the density of calbindin-positive neurons was found to be significantly reduced in the PFC, whereas no such difference was found for parvalbumin-positive cells^91^. Similarly, lowered somatostatin expression was reported in the PFC^24^ and other brain regions^92^ of MDD patients. In contrast, other studies reported decreased PV cell densities or expression levels in the PFC^93^ and other regions including the orbitofrontal and cingulate cortex^91^. Furthermore, changes of perineuronal nets (PNN), a specialized extracellular matrix structure preferentially surrounding parvalbumin-positive cells, were found to correlate with early-life adversity often resulting in depression symptoms during adulthood^94,95^. Parvalbumin and PNNs were even hypothesized to work as predictors of suicidality in humans^16^. PNNs influence PV interneuron activity through multiple mechanisms, such as stabilizing synaptic inputs, modulating their excitability and enhancing the fast-spiking activity^96,97^, supporting the relevance of altered PV interneuron activity for neuropsychiatric disorders.

The crucial role of PV interneurons for MDD was also reproduced in several animal models investigating depression or displaying depression-like symptoms. Both somatostatin and parvalbumin expression in the PFC was dysregulated by stress, leading to depression-like symptoms in mice^20^. Also, region-specific changes in PV cell numbers were reported in depression mouse models, with fewer cells in PFC and hippocampus but increased numbers in the basolateral amygdala^16^. On the other hand, over-inhibition in the hippocampus due to increased *Pvalb* expression and enhanced GABAergic transmission also resulted in depression-like symptoms^17^. While these findings seem contradictory at first glance, they highlight the necessity to correlate the plain cell numbers with the resulting implications for network connectivity and firing rates. In doing so, several studies found diminished PV cell function and disorganized pyramidal cell regulation as functional causes for the expression of symptoms related to depression, schizophrenia, and autism^18,42^. This is in line with our findings, as PV cell numbers were unaltered in all investigated brain regions of *Pvalb-Cre/Dnmt1*-KO mice, while their inhibitory efficacy was greatly diminished (Fig. 1d) due to the DNMT1-mediated effect on GABAergic transmission, which likely causes homeostatic adjustments of network activity (Fig. 1b-c).

Lastly, with epigenetics increasingly emerging as key mechanisms in various disease contexts, it is no surprise that diverse changes in DNA methylation were reported in the context of depression^98–100^. Several disease-relevant genes were highlighted, but investigations in humans were often restricted to blood or saliva samples, leading to findings of increased, decreased or unaltered DNA methylation patterns for the same respective genes^98^. In human blood samples, DNA methylation of the parvalbumin promoter region was found to be significantly increased in several positions, even correlating with disease severity^101^. Supporting this, several studies found DNMT overexpression and downregulation of GAD-genes in GABAergic interneurons of psychosis patients and mice with schizophrenia phenotype^102,103^. While increased promoter methylation could potentially explain the decreased PV expression rates found in some of the aforementioned depression-studies, this inference still needs to be verified.

As neuropsychiatric diseases are complex disorders, the onset, progression and severity of symptoms are concertedly determined by several factors. While changes in cortical E/I balance already constitute a fundamental functional alteration, further consequences of altered gene methylations might also contribute to the observed phenotype. Although most methylation and expression changes in our *Pvalb-Cre/Dnmt1* KO-mouse line are clearly associated to altered synaptic transmitter release (sequencing results published in^21^), there is also a visible overlap of affected genes described in disease-relevant mouse models (see Supp. Tables 1-2). Among them are several strains expressing behavioral traits associated to depression^104–107^ and anxiety^108–110^.

Our observations demonstrate the need for holistic experimental approaches, spanning molecular, neurophysiological, and behavioral experiments to study the role of epigenetic alterations for neuropsychiatric disorders. Using these tools, we found that cortex-wide alterations of PV interneuron function affected the network in a way that still enabled sensory information processing but resulted in depression-like symptoms. As both altered PV neuron function and changes in DNA methylation patterns have been implicated with MDD and we observed a matching behavioral output, our mouse model might be suitable to contribute to the investigation of depression. With DNA methylation having potentially more far-reaching consequences beyond the changed activity of interneurons we reported here, this mouse model bears the potential to shed light on investigating genetic predispositions and flexible disease characteristics such as disease onset and remission likelihood.

## 4 Methods

### 4.1 Animals

Transgenic *Pvalb-Cre/tdTomato/Dnmt1 loxP^2^*-mice were used as knockout animals and *Pvalb-Cre/tdTomato*-mice as controls. Unless stated differently, all experiments were conducted using male mice aged 3–6 months. Both mouse lines were established on a C57BL/6-background by crossing in the *Pvalb-Cre* line (obtained from Christian Huebner, University Hospital Jena, Germany and described in Hippenmeyer et al.^111^) and the *tdTomato* transgenic reporter mouse line (obtained from Christian Hübner, University Hospital Jena, Germany and described in Madisen et al.^112^). To obtain *Dnmt1*-knockout mice, *Dnmt1* floxP^2^ mice (B6;129Sv-Dnmt1^tm4Jae^/J, Jaenisch laboratory, Whitehead Institute; USA; Jackson-Grusby et al.^113^) were crossed in, in which LoxP-sites are flanking exons 4 and 5 of the *Dnmt1* gene. This results in a null *Dnmt1* allele upon Cre-mediated excision^113^. Correct genotypes were validated with the following primers: *Dnmt1* forward 5′-GGGCCAGTTGTGTGACTTGG and reverse 3′-CCTGGGCCTGGATCTTGGGGA (results in a 334 bp wildtype and 368 bp mutant band); *tdTomato* wildtype forward 5′-AAGGGAGCTGCAGTGGAGTA, wildtype reverse 3′-CCGAAAATCTGTGGGAAGTC, mutant forward 5′-CTGTTCCTGTACGGCATGG and mutant reverse 3′-CTGTTCCTGTACGGCATGG (results in a 297 bp wildtype and 196 bp mutant band); *Pvalb*-*Cre* forward 5′-AAACGTTGATGCCGGTGAACGTGC and reverse 3′-TAACATTCTCCCACCGTCAGTACG (resulting in a 214 bp fragment). All animal procedures were performed in strict compliance with the EU directives 86/609/EWG and 2007/526/EG guidelines for animal experiments and were approved by the local governments (Thueringer Landesamt, Bad Langensalza, Germany and Landesamt für Natur, Umwelt und Verbraucherschutz Nordrhein-Westfalen, Recklinghausen, Germany). Unless indicated otherwise, animals were housed under standard housing conditions: mice were kept in compatible groups, under 12 h light/dark conditions, with ad libitum access to food and water, and nestlets and tunnels as environmental enrichments. The weight of individual animals was determined prior to behavioral experiments or prior to sacrificing and brain preparation.

### 4.2 Electrophysiological recordings

#### Surgeries

For all surgeries isoflurane in oxygen was used as anesthesia (3% isoflurane for induction, 1–2% for maintenance), and subcutaneous injections of carprofen (4 mg/kg, Rimadyl, Zoetis GmbH) and buprenorphine (0.1 mg/kg, Buprenovet sine, Bayer Vital GMBH). Eye ointment (Bepanthen, Bayer Vital GmbH) was applied to their eyes to prevent damage from drying out. After medially incising the dorsal skin of the head, it was pushed outward and fixed in place utilizing tissue adhesive (Vetbond, 3M). Dental cement (C&B Metabond, Parkell; Ortho-Jet, Lang Dental) was applied on top of the tissue adhesive to create a support structure for a custom-built circular headbar.

Viral injections (AAV1.shortCAG.dlox.hChR2(H134R).WPRE.hGHp, Zurich Vector Core, viral titer = ∼8x1012 vg/ml) were carried out according to the experimental scheme. In naive mice (Fig. 1a-I, 2a-c), S1 and V1 were targeted unilaterally, whereas in trained animals (Fig. 2g-i, Supp. Fig. 2b-c) bilateral injections in ALM were conducted. The positions of all injections were chosen by stereotactic coordinates (ALM: AP 2.5/ML -1.5; V1: AP -4/ML -2). Injections were conducted using a thin glass pipette with a micropump (Nanoject3, Drummond Scientific) after thinning out the skull with a dental drill. At each site, two quantities of 200 nl were injected at different cortical depths (300/600 µm), adding up to a total volume of 400 nl virus per target region. The 200 nl were injected in 20 steps using 10 nl portions at 14 s intervals. For electrophysiological recordings, circular craniotomies were performed using a biopsy puncher and then covered with a round coverslip of similar size using light-curable dental cement (DE Flowable composite, Nordenta) to fix it in place. The coverslips contained a small opening to later allow access for the Neuropixels probe. Between recordings, this opening was covered with silicon. A thin layer of cyanoacrylate (Zap-A-Gap CA+, Pacer technology) was used to cover the skull. After the surgery, buprenorphine (0.1 mg/kg body weight) was again injected subcutaneously. The animal’s home cage was kept on a heating pad following the surgery until the mouse recovered from anesthesia. To prevent inflammation of the surgical wounds, antibiotics (0.1 ml enrofloxacin/100 ml water) and analgesics (1 ml buprenorphine/100 ml water) were administered via the animal’s drinking water for at least three days. Progression of wound healing and the animal’s general health were scored daily following the surgery.

To record from naive animals, two mice per genotype were utilized; all of them were approx. 6 months of age during surgery. For behavioral analysis and recordings in trained mice, 7 animals (3 control/4 knockout) were used; they were approx. 8 weeks old during surgeries.

#### Neuropixels recordings

Electrophysiological recordings were carried out in head-fixed mice using Neuropixels 1.0 probes. Recordings were carried out on four consecutive days. Probes were painted with DiD cell labeling solution (Invitrogen V22887) before each recording to later identify recording sites in fixed brain slices.

The stimulation protocol was started 5–10 min after insertion and we recorded high pass-filtered data above 300 Hz at 30 kHz and low pass filtered signals between 0.5– 100 Hz at 2.5 kHz from the bottom 384 channels of the Neuropixels probe (∼3.8 mm active recording area). Signals were acquired with an external Neuropixels PXIe card (IMEC, Belgium) used in a PXIe-Chassis (PXIe-1071, National Instruments, USA). Triggers and control signals for different stimuli were separately recorded as analogue and digital signals using the SpikeGLX software (Janelia Farm Research Campus, USA; Bill Karsh, https://github.com/billkarsh/SpikeGLX).

The visual stimulus to test sensory responses in V1 and ALM consisted of a 0.1-s long full-field low-pass filtered Gaussian noise patterns with a cutoff at 0.12 cycles per degree and a temporal frequency of 1 Hz^114^. In each trial, the stimulus was presented twice, with an inter-stimulus interval of 0.5 s. Visual stimulation to induce gamma oscillations consisted of 5-s-long full field drifting square wave gratings at horizontal orientation and a spatial frequency of 0.04 cycles per degree. All visual stimuli were presented at 17-cm distance from the right eye on a gamma-corrected LED backlit LCD monitor (Viewsonic VX3276-2K-MHD-2, 32”, 60Hz refresh rate). The overall stimulation protocol consisted of 50 trials of each type, presented in randomized order with inter-trial intervals between 3–5 seconds.

For optogenetic stimulation, we used a fiber-coupled laser system (ReadyBeam Bio1, FISBA AG), connected to a 105 µm glass fiber (0.22 NA) that was positioned over the cortical recording site. Optogenetic stimulation consisted of a 3-s-long pulse at 488 nm with a peak power of 10 mW from the fiber tip. To avoid transient changes in PV activity, the pulse intensity was first increasing for 1 second, then remained at a peak intensity for 1 second, and lastly decreased back to zero for 1 second. The sequence of visual, optogenetic, and combined trials was fully randomized. For optotagging of PV interneurons, we presented 20 optogenetic stimulus sequences at 5 Hz for 1 second, with an individual pulse length of 10 ms and a peak power of 2 mW from the fiber tip. PV interneurons were then identified by short-latency responses (<2 ms) to individual laser pulses and an increase in firing rate during the 3-s-long optogenetic stimulation period.

To extract spiking activity, channels that were broken or outside of the brain were detected and removed from the recording using the Spike Interface analysis toolbox^115^. The remaining channels were then time-shifted and median-subtracted across channels and time. Corrected channels were then spike-sorted with the Kilosort 2.5 software and manually curated using phy (https://github.com/cortex-lab/phy). All sorted spiking and local field potential data were then analyzed using custom Matlab code (2020b, MathWorks).

Peri-event time histograms (PETHs) for each cluster were computed with a bin size of 10 ms and baseline-subtracted by the mean activity within 1 second before the laser onset. Trial-averaged local field potentials (LFPs) in response to visual stimulation were similarly baseline-corrected. To compute current source densities (CSDs) from LFP signals, we used the inverse CSD method by Pettersen et al.^116^. We applied the spline iCSD method, assuming a smoothly varying CSD between electrode contacts, based on polynomial interpolation. We assumed a homogeneous, isotropic conductivity of σ = 0.3 S/m within and directly above cortex^116^. To reduce spatial noise, the estimated CSD was subsequently convolved with a Gaussian spatial filter with a standard deviation of 0.1 mm. For LFP and CSD analyses, the signals from neighboring contacts at the same depth were averaged together to improve signal quality.

### 4.3 Behavioral Assays

#### Optometry

Assessment of visual acuity and contrast sensitivity were performed as described by Prusky et al.^117^. In brief, each mouse was able to freely turn on a platform surrounded by sine-wave grating stimuli, which varied in spatial frequency and contrast, to create the illusion of a cylindrical stimulus moving around the animal. Intact perception of these stimuli resulted in matching head movements of the mice (optomotor response). To measure the animals’ visual acuity, the spatial frequency (cycles per degree, [cpd]) of stimuli at full contrast (100%) was increased to determine the threshold-value for an optomotor response. Contrast sensitivity of animals was measured by gradually decreasing the stimulus contrast of sinusoidal gratings with distinct spatial frequencies (0.031, 0.064, 0.092, 0.103, 0.192, and 0.272 cpd). The threshold was then calculated as a Michelson contrast from the screen’s luminance ((maximum-minimum)/(maximum+minimum)). Contrast sensitivities for distinct spatial frequencies are displayed as the reciprocal of the threshold. We used 13 control and 11 knockout mice.

#### Touchscreen Chamber

Utilizing operant conditioning, mice were trained to solve a visual contrast discrimination task in a custom-built touchscreen chamber^118^. For this, animals first learned to self-reliantly operate the setup. Therefore, they were water-deprived two days prior to experiments. In the first two basic learning phases, mice were (i) taught to receive a reward in the chamber whenever a green light was turned on and (ii) to touch the screen to receive the reward. Subsequently, a black (full contrast), looming visual stimulus was presented either on the left or right side of the touchscreen. The position of the stimulus on the screen alternated randomly in each trial. Here, animals were required to touch the stimulus to receive a reward. After mice were reliably performing this task, a distractor stimulus with 50% contrast was presented next to the target. Once the mice learned to favor the stimulus with the higher contrast, the distractor’s contrast was randomly varied throughout sessions (Fig. 1i; distractor contrasts: 0%, 50%, 75%, 88.5%). In this 2-alternative forced choice task, we exploited a bias counter-acting mechanism to enforce that mice actively compared the presented stimuli and made sensory-guided decisions to receive a reward. Stimulus-independent strategies, such as repetitively choosing the same side or alternating between them would not yield high reward levels. We used 10 control and 7 knockout mice.

#### Evidence accumulation task

After full recovery from surgeries, animals were water-deprived for one day before introducing them to the behavioral task. To obtain rewards, mice needed to accumulate visual evidence in form of visual grating-like cues and indicate the side with the higher number of cues. Experiments were conducted in a custom-built setup with stimulus presentation, lick-detection and water reward controlled by a microcontroller (Teensy 3.2, PJRC) and Python code (Python 3.7). In general, the task consisted of three periods: 3 s sensory stimulation with randomized sequences of up to 6 cues on one or both sides, 0.5 s delay and a 2-s response window. In between trials was a 3.5-s inter trial interval, which was prolonged after missed trials or incorrect responses. Mice first learned to reliably detect visual stimuli, by indicating the side were all 6 cues (spatial frequency of 0.018 cpd, temporal frequency of 2 Hz) were shown. After mice performed more than 75% of trials correctly in 3 out of the last 4 sessions, distractor stimuli were added on the non-rewarded side. The number of cues on both the target and distractor side varied, resulting in different levels of difficulty depending on the absolute difference in cues shown on either side. Training was conducted using 3 control and 4 knockout animals.

#### Morris water maze (MWM)

Visual navigation and learning capacities were investigated using a MWM. The setup consisted of a cylindric water-filled pool (Ø 1 m) surrounded by an opaque curtain, four distinct landmarks on the pool wall, and a transparent platform (Ø 10 cm). The setup was divided into four virtual quadrants with the platform being consistently in the same quadrant. The start position of the mouse was pseudo-randomly alternated between trials. On the first day of testing, mice started once per quadrant for habituation. The platform slightly protruded from water level to facilitate quicker locating. Upon finding the platform, the animal was required to remain on top of it for 5 s for the trial to be counted successful. If an animal did not find the platform or left it after less than 5 s and did not return onto it within the 90-s period of the trial, the mouse was manually guided to the platform before taking it back to their cages. Between trials, mice were kept under red light for at least 10 min. For the actual learning task, the platform was submerged (1–2 cm) and the maximum duration per trial was set to 60 s. Each mouse started three times per quadrant in a pseudo-random order, equaling 12 trials per day. This training scheme was conducted on five consecutive days. Mice were recorded from above and behavioral data were processed using ANY-maze (Stoelting Co., Wood Dale, USA). We tested 8 control and 8 knockout mice.

#### Home cage activity

To assess the animals’ activity in their home cages, cameras were set up in front of cages of group-housed animals. Each cage, was filmed on several separate days with a framerate of 25 frames per second. From each of these recordings, eight uninterrupted 30 min periods were used for further analysis. For each time period, a pixel-wise comparison with ImageJ was utilized to calculate the difference between every fifth frame of a video. For this, we used the ImageJ Image Calculator and adjusted the threshold to exclude background noise. Subsequently, we measured the area of differing pixels. We filmed 3 control and 3 knockout cages, with 2 mice (6 control/6 knockout mice).

#### Rotarod

A rotarod was used to assess balance and motor performance of mice. Initially, the mice (13 control/11 knockout) were individually habituated for 4 min at a constant speed of 4 rpm. The subsequent test phase comprised three consecutive trials, each lasting a maximum of 5 min, with continuous acceleration from 4 to 40 rpm.

#### Wire hang test

To record neuromuscular strength, mice were placed on a metal wire cage lid, which was then slowly inverted and positioned over a type IV Makrolon cage (Fig. 3d). Each mouse was given three trials of max. 5 min, with inter trial intervals of approx. 3–4 min. In total, 24 mice (13 control/11 knockout) conducted this experiment.

#### Food intake

Food intake of group-housed mice was quantified by assessing the change in cumulative pellet weight after either 24 h or 72 h. Animals remained in their familiar cage settings and pellet weight measurements were calculated relative to the number of animals per cage as the consumed amount per day. Tests were conducted on 4 control and 4 knockout cages (each cage containing 2 mice: 8 control/8 knockout). For each cage two 24-h periods and eight 72-h periods were recorded.

To measure food and water intake in single-housed mice, mice were kept individually for at least two weeks, before measuring pellet and water bottle weight of each cage on five consecutive days. For this, we used 4 control and 6 knockout mice.

#### Nest-building test

Nest-building tests were executed as described by Deacon^119^ and as visualized in Fig. 3f. In brief, mice were weighed and transferred to individual testing cages (IVC cages, Tecniplast) with ad libitum access to food and water, but without any additional environmental enrichment. One nestlet of known weight was placed in each cage. The test was initiated at 5 pm and nest-building skills were scored the next morning at 9 am. Nests were scored according to the following criteria: Score 1 = no noticeable interaction with the nestlet (more than 90% of the nestlet remained intact), Score 2 = nestlet was partially torn (50–90% of the nestlet remained intact), Score 3 = nestlet was largely torn, but no identifiable nest site was build (10–50% of the nestlet remained intact, but less than 90% of shredded material is gathered within a quarter of the cage area), Score 4 = nestlet was used to build a flat nest (less than 10% of the nestlet remained intact, less than 50% of the nest circumference is build higher than the resting mouse), Score 5 = nestlet was used to build a (near) perfect nest (less than 10% of the nestlet remained intact, more than 50% of the nest circumference is build higher than the resting mouse). Each mouse was tested twice with tests being one week apart from each other. In the time between tests, mice remained single-housed without access to further enrichments beside the handed nestlet. Overall, 49 mice (24 control/25 knockout) at ages between 3–6 months were tested.

#### Coat state

The animals’ coat state was assessed in a blinded manner. The scoring system was adapted from Nollet et al.^64^ as visualized in Fig. 3g. Seven different regions of the animal’s body were scored individually with one of three distinct ranks. In case of a good coat state (smooth and shiny fur, no tousled or spiky patches) rank 0 was assigned. Strongly matted, oily or even stained fur was assigned with a rank 1. Slight deviations from the optimum coat state were ranked with a 0.5. Among the investigated areas were both dorsal (head, neck, back, tail) and ventral areas (fore paws, abdomen, hind paws). Overall, 12 control and 12 knockout mice (2–7 months) were investigated.

#### Voluntary wheel running

Single-housed mice were kept in cages with access to running wheels (TSE Systems). Animals were housed for up to three days in these cages. While food and water were provided ad libitum, no further enrichments despite the running wheels were provided. Each animals’ wheel usage was continuously recorded and subsequently evaluated using a custom-written Python script. In total, 11 control and 9 knockout mice were utilized.

#### Interaction test

To test for social interest in female conspecifics, virgin C57BL6J-females were transferred to cages of virgin transgenic, single-housed males. These encounters were filmed for 10 min to evaluate the behavior of the transgenic males. Behaviors such as sniffing, grooming or mounting of the female intruder were evaluated second-wise. The same female encountered two males per day, one per genotype. Between these encounters were at least 6 hours to ensure that the females did not transfer any noticeable odors from the first encounter during the second session of the day. This way we were able to compare the reaction of males of different genotypes to the same female during the same time of her estrus cycle. A total of 16 animals (8 controls/8 knockouts) were utilized.

#### Female urine sniffing test

The reaction of transgenic males towards female olfactory cues was tested using female urine. For this, urine from 5–10 C57BL6J-females from different cages was collected and mixed. To circumvent reactions of the tested males towards the presentation medium, pieces of parafilm were kept in the animals’ cages for one hour prior to testing them. Then, 40 µl tap water were applied to the parafilm as neutral cue, and the time spent sniffing at the application site was assessed for 3 min. After a 45 min break, the same test was conducted using urine instead of water. A total of 16 animals (8 controls/8 knockouts) were utilized.

#### Novelty suppressed feeding test

Experiments were conducted according to Samuels & Hen^120^ and as visualized in Fig. 5a. In brief, mice were weighed and transferred to individual cages (IVC cages, Tecniplast) with environmental enrichments and *ad libitum* access to water. After 24 h of food deprivation, the novelty suppressed feeding test was conducted. For this, mice were individually placed in an arena (i.e., a mouse shipping container with bedding on the ground) with a food pellet attached to a platform in the center. Mice were filmed from above to later analyze their movement patterns using DeepLabCut^121^. Once a mouse started consuming the presented pellet, the animal was transferred to its home cage and a food pellet of known weight was placed in the food tray. After the test, all animals were weighed again. In total, 20 mice (10 control/10 knockout) aged 3–10 months were tested.

### 4.4 Luminex Blood Analysis

To test for hormonal blood concentrations, blood samples were collected *post-mortem*. For each mouse, two samples of 50 µl blood serum were processed. Samples were purified and analyzed using Luminex System (Millipore). Using antibody-coated beads, a multiplex assay was performed to assess changes in hormone levels being quantified using the MAGPIX detection system. The MILLIPLEX Analyst 5.1 software then calculated hormone levels of ghrelin, GLP-1, glucagon, insulin, leptin, PYY, and TNFα in pg/ml. A total of 6 control and 6 knockout mice, all aged 6 months, were used.

### 4.5 Brain Tissue Preparation

Mice were anesthetized using 5% (v/v) isoflurane before decapitation. For immunohistochemistry, mice were transcardially perfused first with phosphate-buffered saline (PBS; pH 7.4), then with 4% paraformaldehyde (PFA) in PBS (pH 7.4). Subsequently, brains were prepared and kept in 4% PFA for 24 h at 4°C for postfixation. Cryoprotection was achieved by incubating the samples in 10% (v/v) and then in 30% (v/v) sucrose in PBS for 24 h each. Brains were subsequently frozen on dry ice and stored at -80°C.

### 4.6 Immunohistochemistry

For immunohistochemistry of free-floating, 40-μm-thick, adult brain sections, slices were first washed three times in PBS/0.05% Triton X-100. GABA-staining required incubation in 1 N HCl for 10 min at 4°C, then in 2 N HCl for 10 min at room temperature and lastly in 2 N HCl for 20 min at 37°C. Afterwards, all sections were placed in citrate buffer (0.05% Tween20, pH 6) for 15 min at 90-100°C. After three washing steps with PBS/0.05% Triton X-100, sections were blocked for 2 h using blocking solution (4% BSA and 10% NGS in PBS/0.05% Triton X-100). Incubation with primary antibodies was conducted for 60 h at 4°C (rabbit anti-GABA (A2052, 1:750), rabbit anti-β-Endorphin (Ab5028, 1:1000), mouse anti-RFP (MA5-15257, 1:500)). Secondary antibodies were applied for 3 h at room temperature (Cy5-donkey anti-rabbit (A10523, 1:1000), Cy3-goat anti-mouse (115035062, 1:1000)). Sections were then stained with DAPI (100 ng/mL in PBS; Sigma-Aldrich) for 5 min. Afterwards, they were mounted onto object slides in pre-warmed 0.5% gelatine supplemented with 0.05% KCrS_2_O_8_ x H_2_O before being embedded in Mowiol (Carl Roth).

### 4.7 Statistics

Statistical tests were carried out using Matlab and Python. Normally distributed data was tested using two-tailed Student’s *t*-test, one-way ANOVA or two-way ANOVA. Non-normally distributed or ordinal data was tested using a Wilcoxon *rank-sum* test. Tukey’s test was conducted as a post-hoc test. The conducted tests for individual experiments are indicated in the text.

## Acknowledgments

We thank Nilufar Nojavan Lahiji and Hendro Langecker for helping with the Morris water maze experiment, Annalena Dobbert for support with the histological analysis, Mira Jakovcevski for methodical assistance in the coat state analysis, Sandra Brill for assistance in the coat state analysis, support in animal breeding/caretaking as well as fruitful scientific discussions.

## Author contributions

J.L.: conceptual design, performed experiments, data analysis, figure illustration, wrote the manuscript; G.N.: method development, experimental supervision, data analysis, figure illustration, manuscript editing; S.G.: performed experiments, data analysis; C.B.Y.: data analysis, manuscript editing; D.P.: experimental supervision, data discussion, manuscript editing; J.R.: performed experiments, manuscript editing; M.H.: performed experiments, manuscript editing; B.K.: data discussion, manuscript editing; A.U.: performed experiments, data analysis, data discussion, manuscript editing; S.M.: performed experiments, data analysis, data discussion, manuscript editing; G.Z.B.: conceptual design, data discussion, wrote the manuscript. All authors have read and agreed to the published version of the manuscript.

