## Supplementary Data for "*Dnmt1*-deficiency in PV interneurons alters cortical circuit function and leads to depression-like behavior"

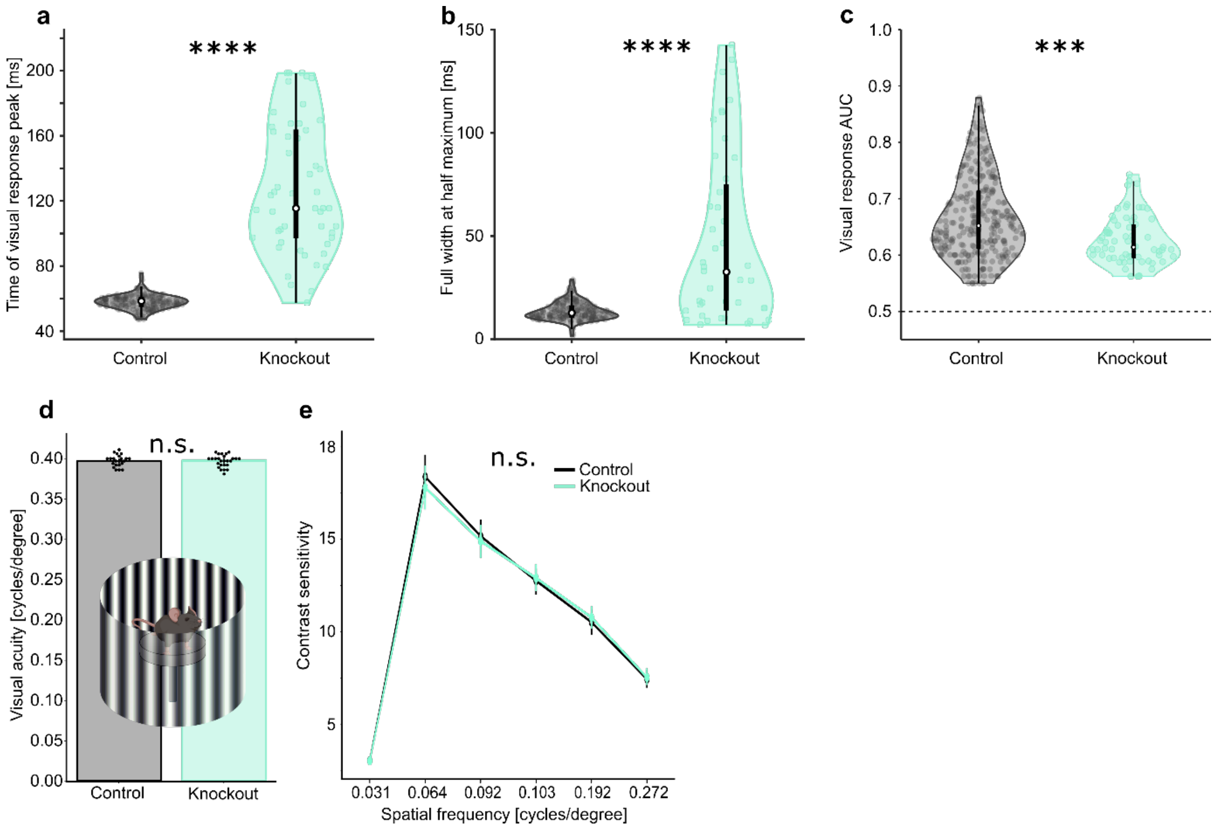


**Supplementary Figure 1: Further quantification of electrophysiological recordings and behavioral outputs.** (**a-c**) Quantification of the traces shown in Figure 1e. (**a**) Time of peak firing rate in response to visual stimulation increased in *Pvalb-Cre/tdTomato/Dnmt1 loxP*$^2$ (knockout) mice compared to *Pvalb-Cre/tdTomato* (control) mice. (**b**) Stimulus responses of mice were less precise as reflected in an increased width at half maximum height. (**c**) The area under the receiver operator characteristic curve (AUC) implies that in response to stimulation, fewer cells were involved in the cortical processing of knockout mice with a lower specificity. (**d-e**) Optometry experiments revealed no behavioral abnormalities in knockout mice ($n$ = 11) in a vision-based test compared to control mice ($n$ = 13). (**d**) The maximum perceived spatial frequency at full stimulus contrast was similar between genotypes. (**e**) Contrast sensitivity in response to stimuli of different spatial frequencies was similar between animals of both genotypes. * $p < 0.5$, ** $p < 0.01$, *** $p < 0.001$, **** $p < 0.0001$.

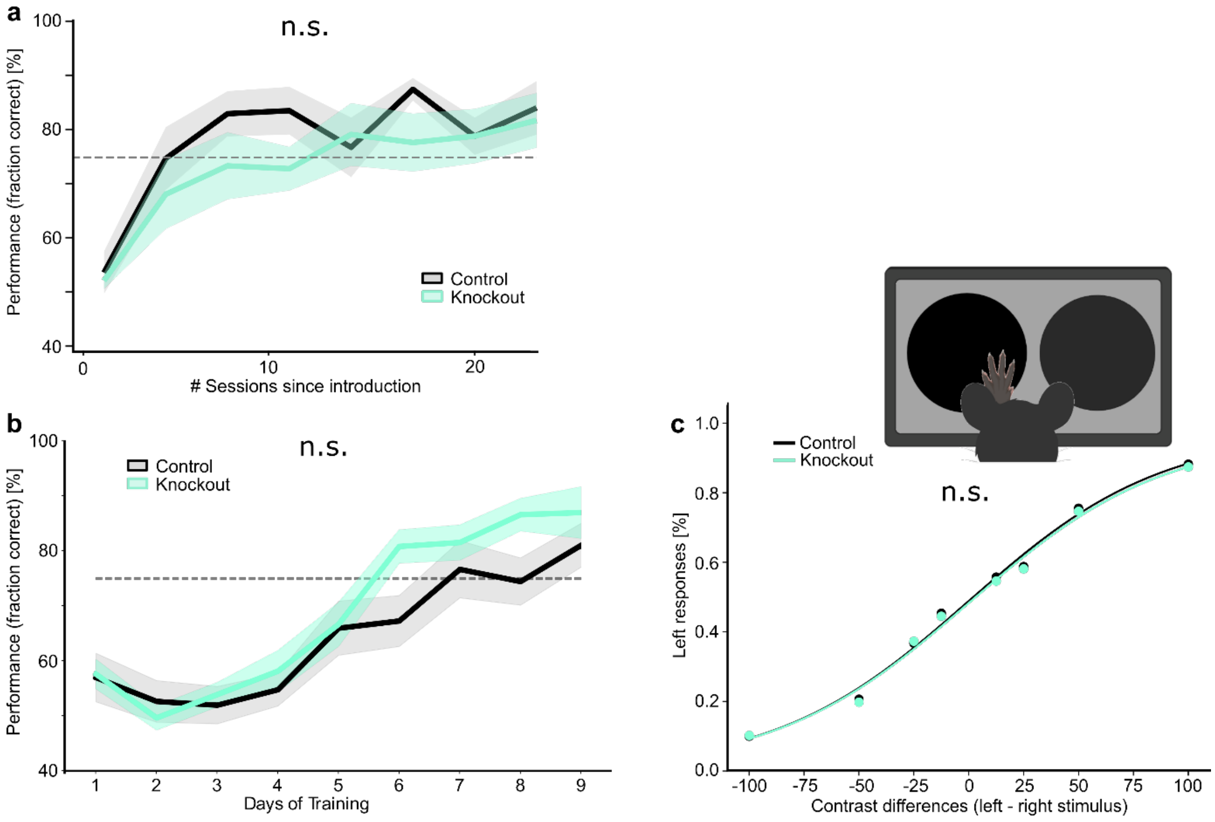


**Supplementary Figure 2: *Pvalb-Cre/tdTomato/Dnmt1 loxP*$^2$ (knockout) mice display intact learning and visual performance compared to *Pvalb-Cre/tdTomato* (control) mice.**
(**a**) Learning curves for a visual evidence accumulation task ($n_{(\text{Control})}$ = 3, $n_{(\text{Knockout})}$ = 4). Corresponding performances are shown in Figure 1j. (**b**) Learning curve and (**c**) performances (mean ± SEM of correct responses) for a contrast discrimination task in a touchscreen chamber ($n_{(\text{Control})}$ = 10, $n_{(\text{Knockout})}$ = 7); n.s. = not significant.

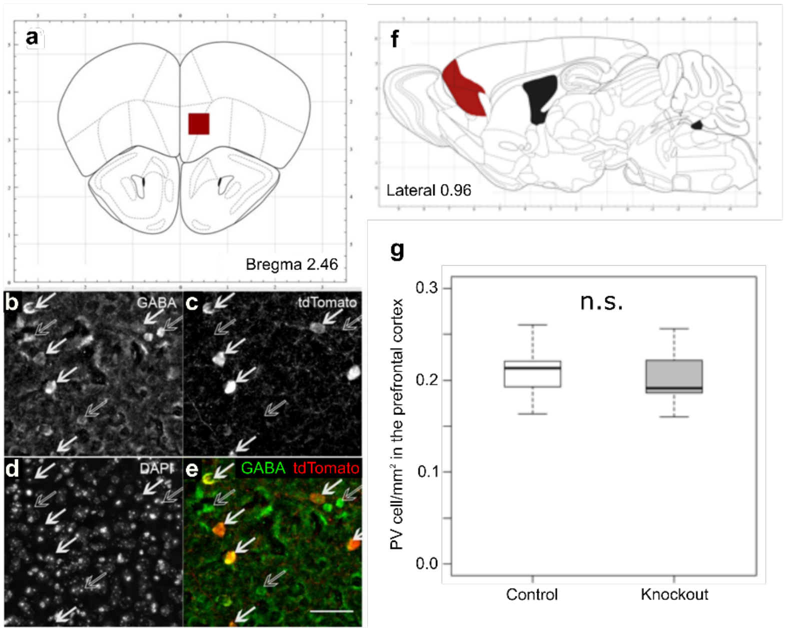


**Supplementary Figure 3: The number of PV interneurons in the prefrontal cortex is not affected by the *Dnmt1* knockout.**

(**a**) Schematic illustration of the prefrontal cortex of a coronally sliced mouse brain at Bregma 2.46. The red square indicates the approximate position of the region depicted in (**b-e**). (**b**) Example image of an immunohistochemical staining against GABA, (**c**) tdTomato, and (**d**) DAPI within the same slice of a 9-month-old *Pvalb-Cre/tdTomato* (control) mouse. (**e**) Overlay of the GABA- and tdTomato-stainings. White arrows show cells with co-expression, while empty arrows indicate GABAergic cells without tdTomato-expression (scale bar = 50 µm). (**f**) Schematic illustration of a sagittally sliced mouse brain with the prefrontal cortex highlighted in red. (**g**) Comparison of PV cell numbers normalized to the area in the prefrontal cortex of *Pvalb-Cre/tdTomato* (control; *n* = 6) and *Pvalb-Cre/tdTomato/Dnmt1 loxP*[2] (knockout; *n* = 7) mice showed no genotype-specific differences.

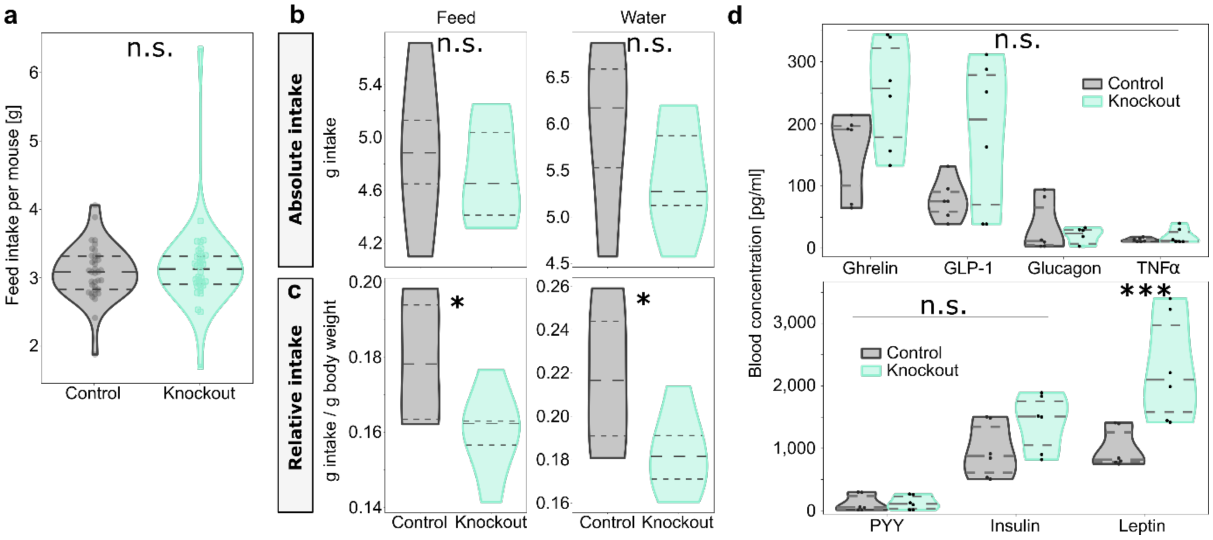


**Supplementary Figure 4: The excess body weight of *Pvalb-Cre/Dnmt1*-knockout mice is likely not appetite-based.**

**(a-c)** Group-housed *Pvalb-Cre/tdTomato* (control) and *Pvalb-Cre/tdTomato/Dnmt1 loxP*[2] (knockout) mice had the same food consumption rates (*n* = 8 mice per genotype). (**b**) In single-housed males ($n_{(Control)}$ = 4, $n_{(Knockout)}$ = 6) both the absolute feed intake as well as the absolute water intake were not altered in *Pvalb-Cre/Dnmt1*-KO mice (upper panels). (**c**) In contrast, the respective intake relative to the animals' body weight showed a significant reduction in food and water intake in knockout mice. (**d**) Blood levels of various appetite-regulating hormones were unaltered by the *Dnmt1*-knockout ($n_{(Control)}$ = 19, $n_{(Knockout)}$ = 16). Prominently, only leptin levels were increased in knockout mice. * $p < 0.5$, ** $p < 0.01$, *** $p < 0.001$, **** $p < 0.0001$.

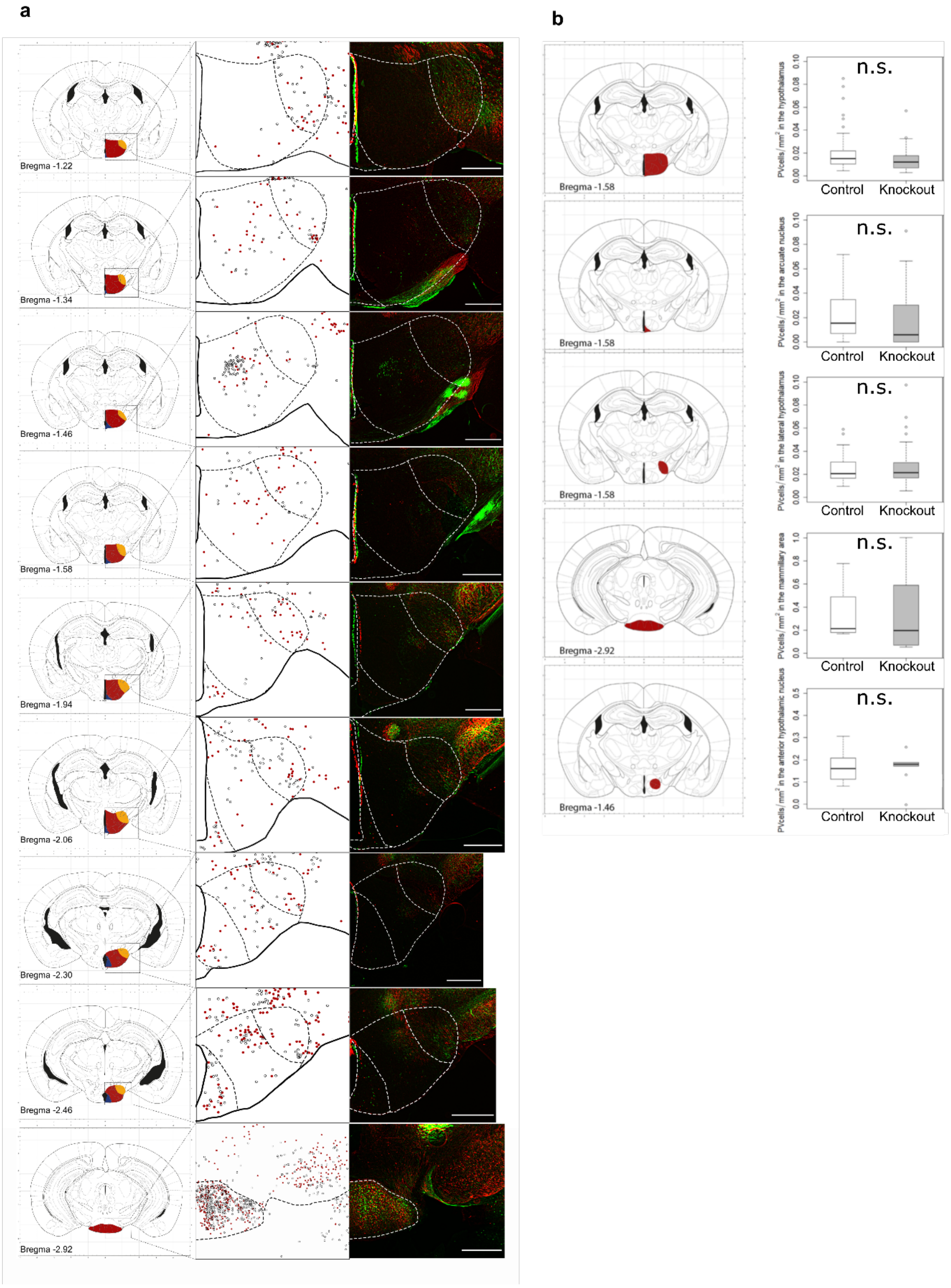


**Supplementary Figure 5: PV cell distribution and quantity were not altered in *Pvalb-Cre/tdTomato/Dnmt1 loxP*$^2$ (knockout) mice**.

(**a**) Cell distribution of tdTomato-labeled PV interneurons was unaffected by the *Dnmt1* knockout. Each row shows a subcortical region relevant in appetite regulation. Left: Schemes visualizing the investigated brain areas. Middle: Overlay of PV cell distribution found in *Pvalb-Cre/tdTomato* (control; grey dots) and knockout (red dots) mice. Right: Pseudocolor overlay of microscopy images of the investigated area. The green color indicates tdTomato-labeling in control mice, red color denotes tdTomato-positive cells in knockout mice. Scale bars = 500 µm. (**b**) Red marked regions in the illustrations on the left-hand side show areas of interest for appetite regulation, where PV cells were counted in slices of different animals ($n_{(Control)}$=4, $n_{(Knockout)}$=3). Quantitative comparisons of PV interneurons in these regions revealed no differences between control and knockout animals (graphs on the right-hand side).

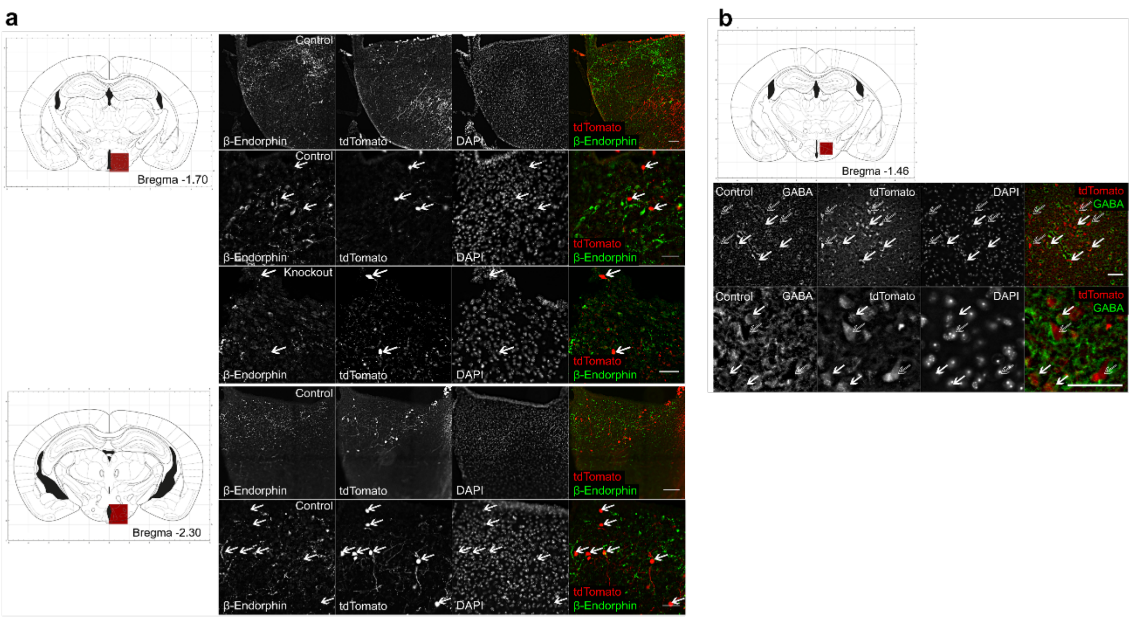


**Supplementary Figure 6: PV cell identity in the hypothalamus is unaltered in *Pvalb-Cre/tdTomato/Dnmt1 loxP*[2] (knockout) mice and does not indicate a connection between PV interneuron activity and appetite regulation.**
(**a**) β-Endorphin immunostaining in the arcuate nucleus of the hypothalamus was conducted to identify Proopiomelanocortin-expressing (POMC) cells. The tdTomato-labeling (white arrows) did not overlap with the β-endorphin staining in both *Pvalb-Cre/tdTomato* (control) and knockout mice (scale bar = 50 µm). (**b**) GABA immunostaining in the anterior hypothalamic nucleus of a control mouse showed a partial overlap of tdTomato- and GABA-labeling (white arrows). Immunostainings indicated that only a fraction of the tdTomato-positive PV interneurons in the hypothalamus express GABA and could thus be attributed to the group of AgRP-expressing cells, while other PV cells were not positive for GABA (empty arrows; scale bar = 50 µm).

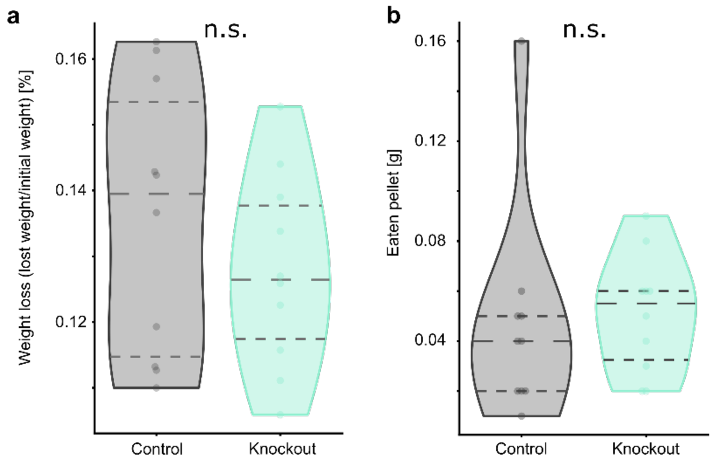


**Supplementary Figure 7: Food deprivation in the novelty-suppressed feeding test affected mice of both genotypes similarly.**
(**a**) *Pvalb-Cre/tdTomato/Dnmt1 loxP*[2] (knockout) and *Pvalb-Cre/tdTomato* (control) mice (*n* = 10 mice per genotype) lost similar amounts of weight after food deprivation. (**b**) In their home cages, mice of both genotypes consumed similar amounts of the pellet in a 5-min time window.

**Supplementary Table 1: *Dnmt1*-knockout in PV interneurons induces upregulated expression of genes also affected in other mouse models.**

Mouse lines marked with an asterisk have been found to express behavioral alterations associated with depression and/or anxiety.

| Mouse line | Enrichment FDR | Number of Genes | Fold Enrichment |
|---|---|---|---|
| ***Htt* Mutation (HTT exon1 142Q) mouse*** | **2.70E-06** | **21** | **5** |
| *Pmp22* KO mouse | 9.90E-05 | 16 | 4.9 |
| *Pmp22* OE mouse | 3.20E-04 | 15 | 4.7 |
| *Pmp22* KO mouse | 1.00E-03 | 13 | 4.3 |
| *Ache AChE-S* mutation mouse | 3.30E-04 | 16 | 4.3 |
| ***Htt* Mutation (HTT exon1 18Q) mouse*** | **9.00E-04** | **14** | **4.1** |
| *Dysf* Deficiency mouse | 1.50E-03 | 13 | 4.1 |
| *Nfatc3* OE mouse | 1.00E-03 | 14 | 4.1 |
| *Tlr4* mutation mouse | 6.30E-05 | 21 | 4 |
| *Emx2* Deficiency mouse | 8.90E-04 | 15 | 4 |
| ***Dmd* Deficiency mouse*** | **3.50E-04** | **17** | **4** |
| ***Rps6ka3* knockout mouse*** | **3.60E-04** | **17** | **3.9** |
| ***Ache AChE-R* overexpressor mutation mouse*** | **9.00E-04** | **15** | **3.9** |
| *Sox9* OE mouse | 8.90E-04 | 16 | 3.8 |
| *Hras* mouse | 1.20E-03 | 15 | 3.8 |
| *CHRNA9* KO mouse | 8.90E-04 | 16 | 3.7 |
| *Atxn7* OE mouse | 3.30E-04 | 19 | 3.7 |
| *Prmt5* KO mouse | 1.60E-03 | 15 | 3.6 |
| ***Sod1 G93A* mutation mouse*** | **8.90E-04** | **19** | **3.3** |
| ***Dicer1* KO mouse*** | **1.90E-03** | **17** | **3.2** |

**Supplementary Table 2: *Dnmt1*-knockout in PV interneurons induces downregulated expression of genes also affected in other mouse models.**

Mouse lines marked with an asterisk have been found to express behavioral alterations associated with depression and/or anxiety.

| Mouse line | Enrichment FDR | Number of Genes | Fold Enrichment |
|---|---|---|---|
| *Prmt1* KD mouse | 1.50E-23 | 51 | 6.5 |
| *Cyfip1* OE mouse | 4.90E-05 | 14 | 5.5 |
| ***Htt* Gain of function by knock in of htt cag repeat length-7/16 mouse*** | **4.90E-05** | **14** | **5.4** |
| *Prmt8* KD mouse | 1.30E-12 | 35 | 5.2 |
| *Eomes* KO mouse | 1.90E-07 | 22 | 5.2 |
| *Nrl* KO mouse | 5.70E-07 | 21 | 5 |
| *Top2b* KO mouse | 4.90E-05 | 16 | 4.7 |
| *Pten* KO mouse | 4.90E-06 | 20 | 4.5 |
| ***Ppard* KO mouse*** | **1.40E-04** | **15** | **4.4** |
| *FOXO3* KO mouse | 1.30E-06 | 23 | 4.3 |
| *Apod* KO mouse | 4.90E-06 | 21 | 4.3 |
| ***Htt* OE mouse*** | **4.90E-05** | **18** | **4.1** |
| ***Setdb1* KO mouse*** | **4.90E-05** | **19** | **4** |
| *Nrl* Deficiency mouse | 1.40E-04 | 17 | 4 |
| ***Atxn1* Knock-in mouse*** | **1.60E-04** | **17** | **3.9** |
| *Mef2d* KD mouse | 4.90E-05 | 20 | 3.8 |
| *Nrl* Deficiency mouse | 6.40E-05 | 20 | 3.7 |
| ***Htt* Knock-in mouse*** | **1.20E-04** | **19** | **3.7** |
| ***Cstb* KO mouse*** | **7.60E-05** | **21** | **3.5** |
| ***Dicer1* KO mouse*** | **1.60E-04** | **20** | **3.4** |
